# Cell cycle-dependent mRNA localization in P-bodies

**DOI:** 10.1101/2024.04.16.589748

**Authors:** Adham Safieddine, Marie-Noëlle Benassy, Thomas Bonte, Floric Slimani, Oriane Pourcelot, Michel Kress, Michèle Ernoult-Lange, Maïté Courel, Emeline Coleno, Arthur Imbert, Antoine Laine, Annie Munier Godebert, Angelique Vinit, Corinne Blugeon, Guillaume Chevreux, Daniel Gautheret, Thomas Walter, Edouard Bertrand, Marianne Bénard, Dominique Weil

## Abstract

Understanding the dynamics of RNA targeting to membraneless organelles is essential to disentangle their functions. Here, we investigate how P-bodies (PBs) evolve during cell cycle progression. PB purification across the cell cycle uncovers widespread changes in their RNA content, which are partly uncoupled from cell cycle-dependent changes in RNA expression. Single molecule FISH shows various mRNA localization patterns in PBs peaking in G1, S, or G2, with examples illustrating the timely capture of mRNAs in PBs when their encoded protein becomes dispensable. Yet, rather than directly reflecting absence of translation, cyclic mRNA localization in PBs can be controlled by RBPs, such as HuR in G2, and by RNA features. Indeed, while PB mRNAs are AU-rich at all cell cycle phases, they are specifically longer in G1, possibly related to post-mitotic PB reassembly. Altogether, our study supports a model where PBs are more than a default location for excess untranslated mRNAs.

## Introduction

Proper compartmentalization of biological molecules is a fundamental aspect of cellular organization. Recently, membraneless organelles received increasing attention for their contribution in organizing subcellular space^1–5^.

P-bodies (PBs) are ribonucleoprotein granules (RNPs) widespread throughout eukaryotes and constitutively present in mammalian cells^6^. They contain a variety of RNA binding proteins involved in translation regulation and RNA decay. Purification of PBs from asynchronous human cells previously allowed the characterization of their RNA content^7^. PBs accumulate one third of the coding transcriptome. These mRNAs are generally abundant, inefficiently translated and strikingly AU-rich^8^. This nucleotide bias in the coding sequence (CDS) results in a low-usage codon bias and poor protein yield. In the 3’ untranslated regions (3’UTRs) it favors accumulation in RNP granules, likely due to the binding of condensation-prone RNA-binding proteins (RBP). PB-enriched mRNAs tend to encode regulatory proteins whereas the mRNAs encoding house-keeping proteins tend to be excluded from PBs^7^. While these studies provided a first picture of the PB transcriptome, further studies are required to understand to which extent PBs adapt their content to cellular needs.

Various approaches have been used to investigate RNA localization in PBs. In situ hybridization first showed that mRNA repressed by miRNA localize to PBs^9^. Then, single molecule tracking of microinjected RNAs allowed examining their trafficking at a high temporal resolution over minutes^10^. This revealed that miRNAs, repressed mRNAs, and lncRNAs can associate transiently or stably with PBs. Other studies addressed RNA localization in PBs in response to translational stress. For instance, relief of miR122-mediated silencing in response to amino acid starvation or oxidative stress causes the release of its CAT-1 mRNA target from PBs^11^. Amino acid starvation leading to 60% translation reduction, also increased exogenous RNA targeting to PBs^12^. While such studies suggested a regulatory potential of mRNA targeting to PBs under conditions of extreme translational reprogramming, the compositional dynamics of PBs in unstressed conditions was not investigated.

The cell cycle is a program of physiological and molecular changes a cell undergoes to produce two daughter cells. From an RNA metabolism perspective, cell cycle progression involves specific waves of transcription^13–18^ and degradation^19–23^. In terms of translation, proliferative cells have a distinct tRNA signature compared to differentiating or arrested cells, which may favor the translation of mRNAs with AU-rich codon usage^24^. Regarding PBs, it was shown that they dissolve at every mitosis, reform in G1 phase, and enlarge during S phase progression^25^. Interestingly, we previously found that, among the various regulatory proteins encoded by PB mRNAs, cell cycle regulators were particularly enriched^6,7^. We thus chose to investigate the dynamics of RNA localization in PBs across the cell cycle.

Analyzing the composition of PBs purified at various cell cycle stages revealed widespread changes in their RNA content during cell cycle progression. Some changes were uncoupled from those occurring in the cytoplasm (due to variations in mRNA transcription or stability), demonstrating a regulatory potential of PBs. Single molecule FISH (smFISH) confirmed the diversity of PB localization patterns with some G2-induced mRNAs trafficking to PBs specifically in early G1, when their encoded protein is no longer needed. Puromycin experiments showed that the cyclic pattern of PB localization is not directly related to the amount of untranslated transcripts in the cytoplasm. Rather, preventing HuR-mRNA interactions abolished cyclic mRNA localization in PBs, particularly in G2. While we found no evidence of phase-specific nucleotide or codon bias, we observed a marked bias in mRNA length in G1. Altogether, our results demonstrate the existence of controlled differential RNA localization in PBs across the cell cycle.

## Results

### PBs enlarge during cell cycle progression

Before analyzing the content of the PBs during the cell cycle, we first refined the description of their global morphology and number across the cell cycle. To identify cell cycle stages, we relied on the PIP-FUCCI system (see Methods, Figure S1A, B). PB labeling by immunofluorescence (IF) against the classical PB marker DDX6 (Figure 1A) showed that PB size increased from G1 to G2 (Figure S1C) and DDX6 intensity in PBs increased in G2 (Figure S1D) regardless of PB size (Figure 1B). However, the number of PBs per cell increased only modestly (Figure S1E). As these changes could result from increased expression of PB proteins required for (DDX6, LSM14A and 4E-T) or contributing to (PAT1B) human PB assembly^26^, we separated HEK293- FUCCI cells in G1, S and G2/M phases using fluorescence activated cell sorting (FACS). However, none of the tested proteins was induced across the cell cycle (Figure S1F). In summary, after dissolving in mitosis, PBs form in G1, progressively enlarge during interphase, particularly in G2, while their number remains similar.

**Figure 1:**
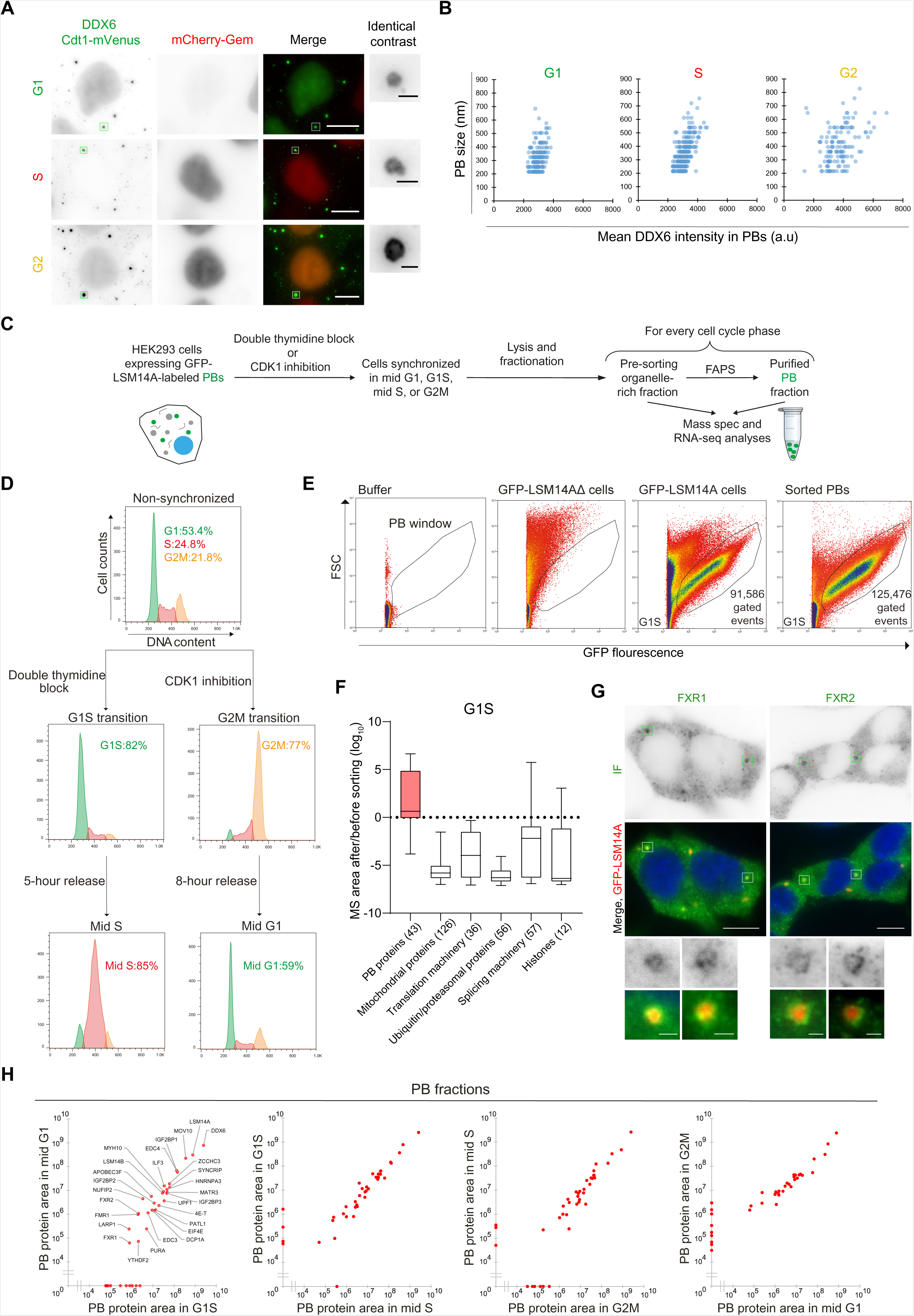
PBs enlarge during cell cycle progression while maintaining a similar proteome. (A) HEK-FUCCI cells with DDX6-labeled PBs. Anti-DDX6 IF reveals cytoplasmic PBs while nuclear Cdt1-mVenus signal indicates cells in G1 or G2 phase (both in green in the merge). Nuclear mCherry-Gem signal indicates cells in S or G2 phase (red in the merge). (B) Scatter plot of PB size and DDX6 intensity across the cell cycle (2 independent experiments). (C) Main steps of PB purification. (D) Cell cycle analysis from synchronized samples, with cells in G1, S and G2M colored in green, red and yellow, respectively. (E) FAPS profiles of pre-sorting lysate and sorted PBs from cells synchronized at the G1S transition. (F) Enrichment or depletion of several protein groups after sorting (ratio of MS area after/before sorting) in cells at the G1S transition. The number of detected proteins (MS area >1) is indicated in brackets. (G) IF of FXR1 and FXR2 in HEK293-GFP-LSM14A cells. The IF and PB signals are green and red, respectively. (H) Pairwise comparison of the abundance (MS area) of known PB proteins (list in Table S1) in the PB fraction between successive cell cycle stages. Scale bars: 10 µm in panels, 1 µm in insets.

### PB purification reveals the same major PB proteins across the cell cycle

To characterize changes in PB composition during cell cycle progression, we purified them from different phases of the cell cycle using our previously developed Fluorescence Activated Particle Sorting (FAPS) method^7^ (Figure 1C) and HEK293-GFP-LSM14A cells. In these cells, GFP-LSM14A was expressed at levels similar to endogenous LSM14A and co-localized with the PB marker DDX6 but not the stress granule marker TIA1 (Figure S1G-J). Cells were synchronized in mid G1, G1S transition, mid S and G2M transition using a double thymidine block or the selective CDK1 inhibitor^27^ RO-3306 (see Methods, Figure 1D). Forty 15 cm plates were synchronized per phase to obtain sufficient material for FAPS purification. Cytoplasmic lysates were then prepared, with an aliquot kept aside for further analysis, termed the pre-sorting fraction (PSF), while the remaining was used to sort PBs (see Methods, Figures 1E and S2A). After sorting, a fraction of each purified sample was re-analyzed by flow cytometry and visualized under microscope to verify that PBs were conserved while contaminants were considerably reduced (Figures 1E and S2B).

After a total of >140 hr sorting, sufficient material was accumulated for proteomic analysis of PBs from each cell cycle phase, using liquid chromatography-tandem mass spectrometry (LC-MS-MS, Table S1). In all phases, the vast majority of known PB proteins (28- 43 proteins detected) were enriched in purified PBs compared to PSFs (Figures 1F, S2C-E).

These included translational repressors and RNA decay factors (DDX6, LSM14A, LSM14B, 4E- T, EDC4 and DCP1A; a tentative list combining established PB proteins and FAPS-suggested candidates is proposed in Table S1). In contrast, proteins such as mitochondrial proteins, translation initiation factors, proteasome subunits, splicing factors and histones were mostly depleted (Figures 1F, S2C-E, Table S1). The presence of FXR1 and FXR2 in PBs was confirmed by IF, which revealed a particular crown-like localization around PBs (Figure 1G). Given the inherently limited amounts of proteins obtained from purified PBs, we could not carry out the replicates needed for quantitative MS. Nevertheless, a pairwise comparison of PBs from successive cell cycle phases did not show drastic changes in the levels of the well-detected PB proteins (Figure 1H). In summary, the proteomic analysis of purified PBs does not reveal major changes of the main PB proteins across the cell cycle.

### PB RNA content is dynamic during cell cycle progression

We then analyzed the RNA content of the purified PBs and their corresponding PSFs using RNA sequencing. First, we assessed if the PSFs recapitulated known cell cycle-dependent RNA regulation. Most RNAs previously reported as more expressed in G1S or in G2M in HeLa cells^28^ were also more abundant in our G1S or G2M PSFs, respectively, despite the different cell line and protocols (Figure S3A). Moreover, gene ontology (GO) analysis showed that RNAs more abundant in our G2M sample encode proteins involved in chromatin condensation, chromosome segregation, and cell division, while those more abundant in G1S encode proteins involved in nucleotide metabolism, DNA replication and its regulation (Figure S3B,C). Altogether, these analyses confirmed the reliability of our datasets. We then compared RNA levels before and after sorting within each cell cycle phase (Figures 2A and S3D). Between 3,838 and 5,106 mRNAs were enriched in PBs (p-adj<0.05, DEseq2 model^29^) depending on the phase (Table S2). Among them, 2043 were enriched in only a subset or a unique phase (Figure 2B).

**Figure 2:**
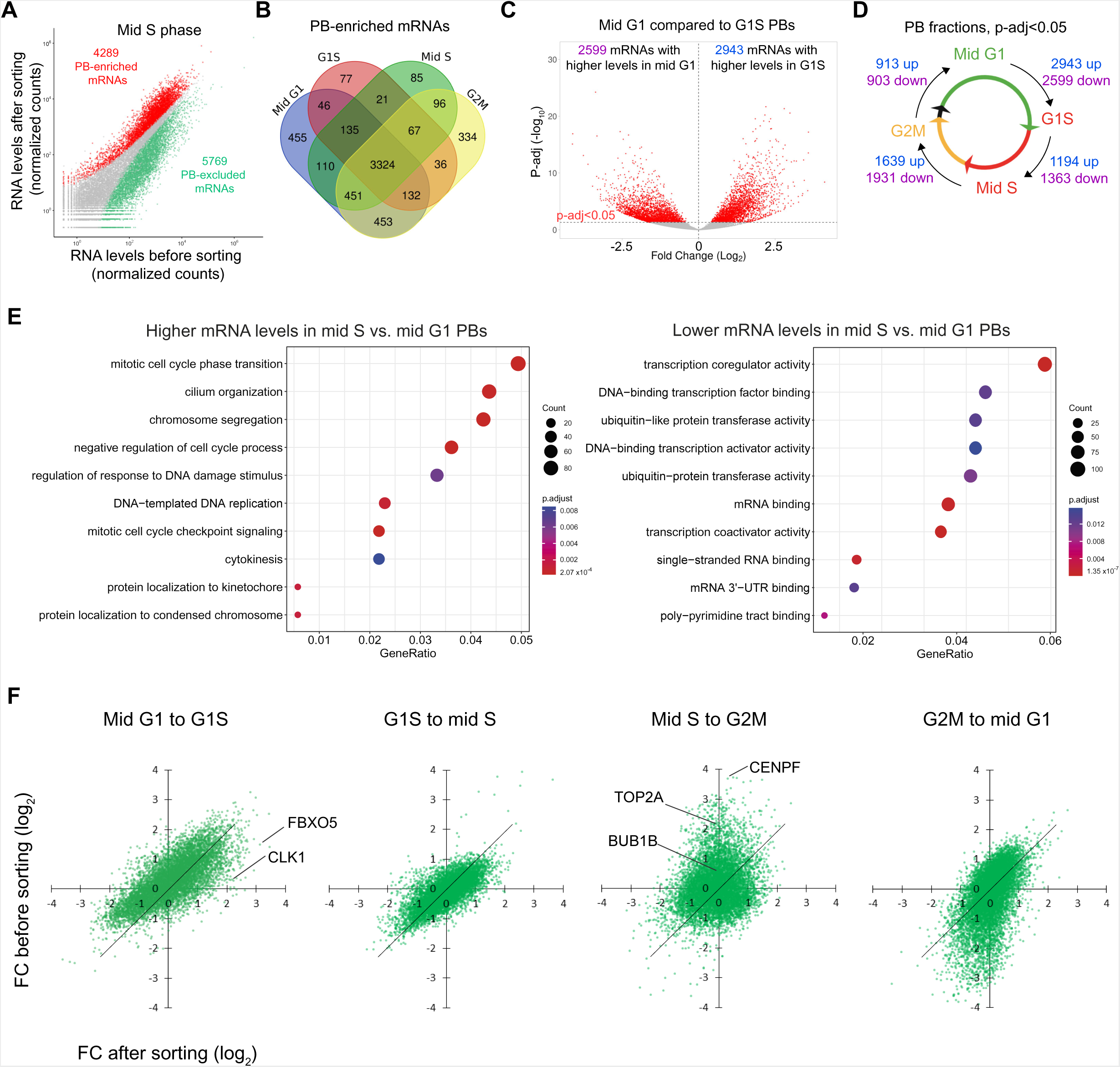
Analysis of purified PBs uncovers their dynamic transcriptome. (A) RNA levels (normalized DEseq2 counts) before and after PB sorting from mid S cells. All expressed mRNAs are shown. The mRNAs significantly enriched and depleted in PBs (p-adj<0.05) are in red and green, respectively. (B) Venn diagram of PB-enriched mRNAs in the different cell cycle phases. (C) Volcano plot showing the changes in PB mRNA content from mid G1 to G1S. All expressed mRNAs are shown with significant changes (p-adj<0.05) in red. (D) Summary of significant changes in PBs (p-adj<0.05) across the cell cycle. (E) GO analyses of PB mRNAs with significant positive (left panel) and negative (right panel) changes in mid S compared to mid G1. The 10 most enriched GO categories are shown. (F) Quadrant plots comparing mRNA fold-changes before and after sorting (normalized counts >50) between successive cell cycle stages. mRNAs further studied by smFISH are indicated. Associated volcano plots are in Figure S4B-D.

Next, we compared the RNA content of purified PBs across the cell cycle (Table S3 and S4, Figure S4A). Remarkably, this revealed widespread changes in RNA levels in PBs (Figures 2C, D and S4B). For example, from mid G1 to G1S, 2943 mRNAs had increased levels in PBs, while 2599 decreased. To determine if the mRNAs with cyclic accumulation in PBs encode proteins enriched for specific functions, we systematically performed GO analyses (Figure 2E). We found that mRNAs accumulating in PBs between mid G1 and mid S were enriched (181 out of 1889) for transcripts encoding proteins functioning in mitotic cell cycle phase transition, chromosome condensation, and cytokinesis. Inversely, the mRNAs that decreased in PBs between mid G1 and mid S were enriched (154 out of 1813) for transcripts encoding proteins involved in transcription, transcriptional regulation, and RNA binding. This indicated that the mRNA content of PBs is highly dynamic throughout the cell cycle.

### RNA accumulation in PBs is partly uncoupled from their cytoplasmic expression level

Cell cycle variations in mRNA levels are controlled at the levels of transcription and/or stability^13,17–20,22,23,28^. The question here was whether there is an additional regulation at the level of mRNA localization in PBs. If not, mRNA levels in PBs should follow their level in the cytoplasm across phases. We therefore investigated the relationship between the RNA content of PBs and the surrounding cytoplasm, using PSFs as a proxy. Comparing RNA changes in PBs and PSFs from mid G1 to G1S showed an overall distribution along the diagonal (Figure 2F, S4B-D), suggesting a broad coupling between the PB and cytoplasm contents. Yet mRNAs decreasing in PBs tended to decrease twice less in the cytoplasm and some cases fell far from the general distribution, as exemplified by FBXO5 and CLK1 mRNAs (studied later), indicating some degree of uncoupling between the PB and cytoplasm contents. The pattern was similar from G1S to mid S. In contrast, from mid S to G2M and G2M to mid G1, the PB content was weakly coupled to the prominent cytoplasmic up and down regulations. For example, it did not mirror the cytoplasmic increase in TOP2A, BUB1B, and CENPF levels (Figure 2F, studied later). These data therefore suggested that the dynamics of the PB transcriptome can be uncoupled from cytoplasmic variations, particularly during the second half of the cell cycle.

### *In-situ* confirmation of various patterns of cyclic RNA localization in PBs

To support our transcriptomic analysis, we next analyzed several mRNAs (chosen based on PB-enrichment pattern, p-adj values, and expression levels for reliable detection) using smFISH, which allows *in-situ* localization and counting of individual RNA molecules^30,31^.

We started with the mRNA of FBXO5, a major regulator of the anaphase promoting complex (APC)^32–37^. In our RNA-seq dataset, FBXO5 expression was highest in G1S and mid S phases in the PSF, in accordance with its reported transcriptional induction at G1S^38^. In purified PBs, it also peaked at G1S and mid S, but with a much higher differential than mid G1 or G2 (Figure 3A). For smFISH, we synchronized HEK293-GFP-LSM14A cells as performed for PB purification. In agreement with the RNA-seq data, we observed more FBXO5 mRNA molecules in the cytoplasm, and considerably more in PBs, in G1S and mid S than in G2M and mid G1 (Figure 3B, C).

**Figure 3:**
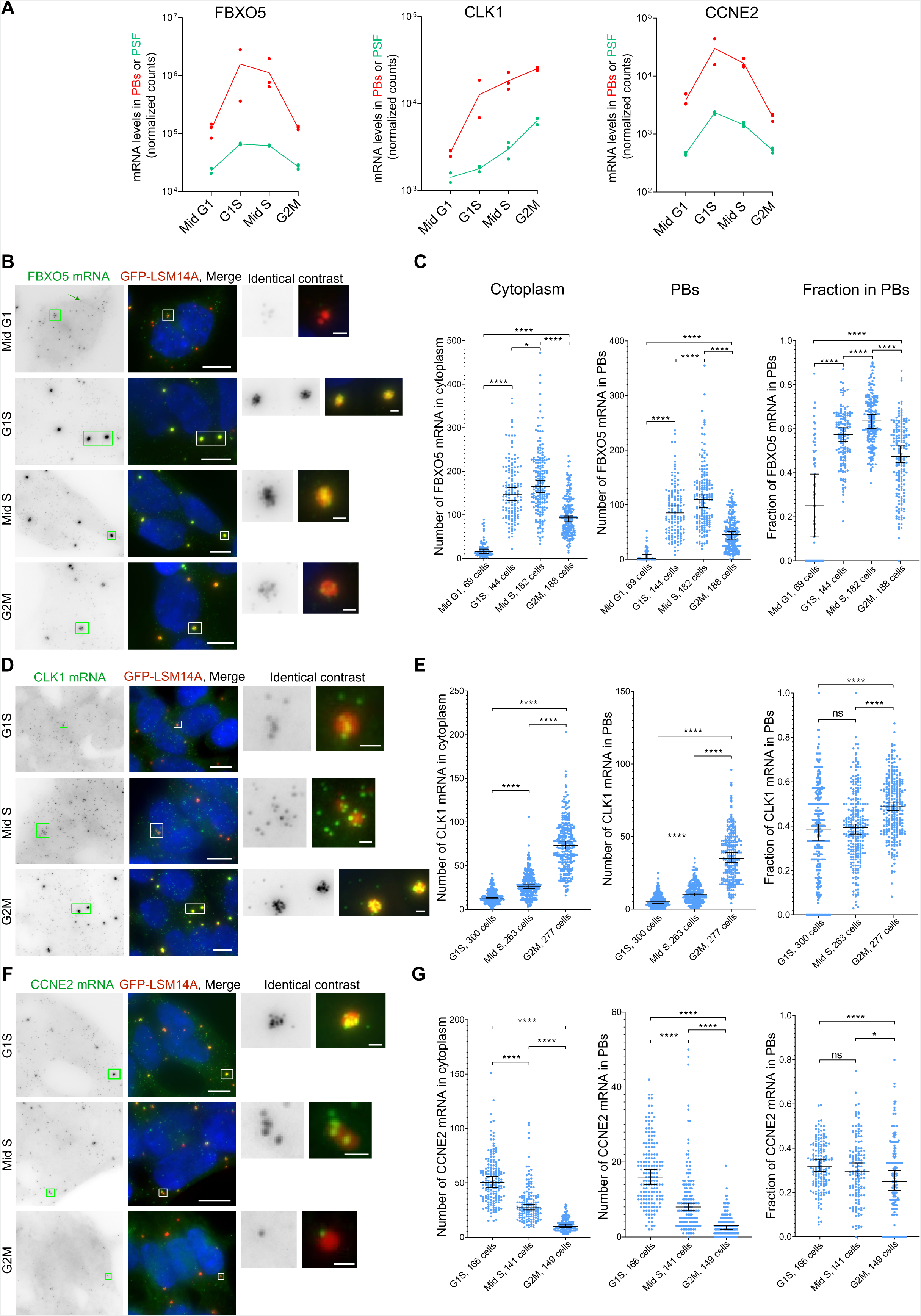
mRNA localization in PBs can peak at any cell cycle phase. (A) Evolution of FBXO5, CLK1 and CCNE2 mRNA levels (in normalized counts) in purified PBs and pre-sorting fractions across the cell cycle. Each dot corresponds to one replicate, with the line connecting their mean. (B) Synchronized HEK293-GFP-LSM14A cells after smFISH of FBXO5 mRNA. For better visualization, the Cy3 smFISH signal is in green and GFP-LSM14A in red. Nuclei were stained with DAPI (blue). Individual mRNA molecules appear as small dots (indicated by the arrow). Insets present enlargements of representative PBs, illustrating the heterogeneity of RNA amounts in PBs. Scale bars: 10 µm in panels, 1 µm in insets. (C) Number of FBXO5 mRNA molecules in the cytoplasm or in PBs, and the fraction of mRNA in PBs across the cell cycle. Each dot corresponds to one cell (2 independent experiments). Horizontal lines, median; error bars, 95% CI. Two-tailed Mann-Whitney statistical tests: ****, p<0.0001; *, p<0.05; ns, non-significant (p>0.05). (D, E) Same as in B and C, for CLK1 mRNA. (F, G) Same as in B and C, for CCNE2 mRNA.

Similarly, we examined the mRNAs encoding the cell cycle-regulated splicing factor CLK1^39^ and cyclin E2 (CCNE2). In contrast to FBXO5, CLK1 transcripts gradually increased in both PBs and the cytoplasm throughout the cell cycle, peaking at G2M (Figures 3A, D, E). CCNE2 transcripts had a similar profile as FBXO5, with both sequencing data and smFISH experiments showing maximal cytoplasmic and PB levels in G1S (Figure 3A, F, G). Thus, CCNE2, FBXO5 and CLK1 mRNAs exhibited various PB localization patterns, peaking at G1S, mid S, and G2M, respectively. These experiments provided single-molecule evidence of cyclic mRNA accumulation in PBs.

### PBs capture a subset of cyclic mRNAs after mitotic exit

We next focused on mRNAs harboring prominent cytoplasmic induction in G2 with limited consequences on PB accumulation (Figure 2F). We first analyzed the mRNA encoding the G2 protein topoisomerase 2A (TOP2A), which regulates chromosome compaction^40^ and relieves topological DNA stress^41,42^. In the sequencing dataset, expression of TOP2A peaked in G2M, as previously described^43–45^. This increase however was not mirrored in PBs (Figure 4A). SmFISH experiments on asynchronous cells revealed three TOP2A localization patterns (Figure 4B): (i) few TOP2A mRNA molecules in both the cytoplasm and PBs; (ii) high TOP2A mRNA expression in the cytoplasm but modest localization in PBs; and unexpectedly (iii) in a subset of cells, heavy TOP2A mRNA accumulation in PBs with few TOP2A mRNA molecules free in the cytoplasm. Quantitative image analysis further assessed this striking heterogeneity (Figure 4C). TOP2A biology^46^ and our RNA-seq data suggested that cells with high RNA expression and low PB localization (pattern ii) corresponded to G2 cells, where the protein is needed and most highly expressed. We speculated that after mitosis, in early G1 (a time point not included in the PB purification pipeline, but accessible with smFISH), leftover TOP2A mRNA could be sent to PBs (pattern iii). To test this hypothesis, we transiently expressed Halo-tagged LSM14A in HEK293-FUCCI cells to allow for the simultaneous imaging of cytoplasmic TOP2A mRNAs, PBs and nuclear PIP-FUCCI markers (Figure 4D). As expected, S and G2 cells showed high levels of TOP2A mRNA with modest PB localization. In contrast, G1 cells showed two patterns, with either minute or moderate TOP2A expression but striking accumulation in PBs. Since TOP2A remains abundant during mitosis^45^ (Figure 4D) and is transcriptionally induced only in G1S or early S^47,48^, we conclude that the latter cells correspond to early G1 where residual TOP2A mRNAs are localized in PBs. To summarize, the burst of TOP2A mRNA expression in G2 cells does not coincide with PB accumulation. Rather, localization in PBs occurs when they reform in early G1, at a time when TOP2A mRNA is down-regulated and the protein no longer required.

**Figure 4:**
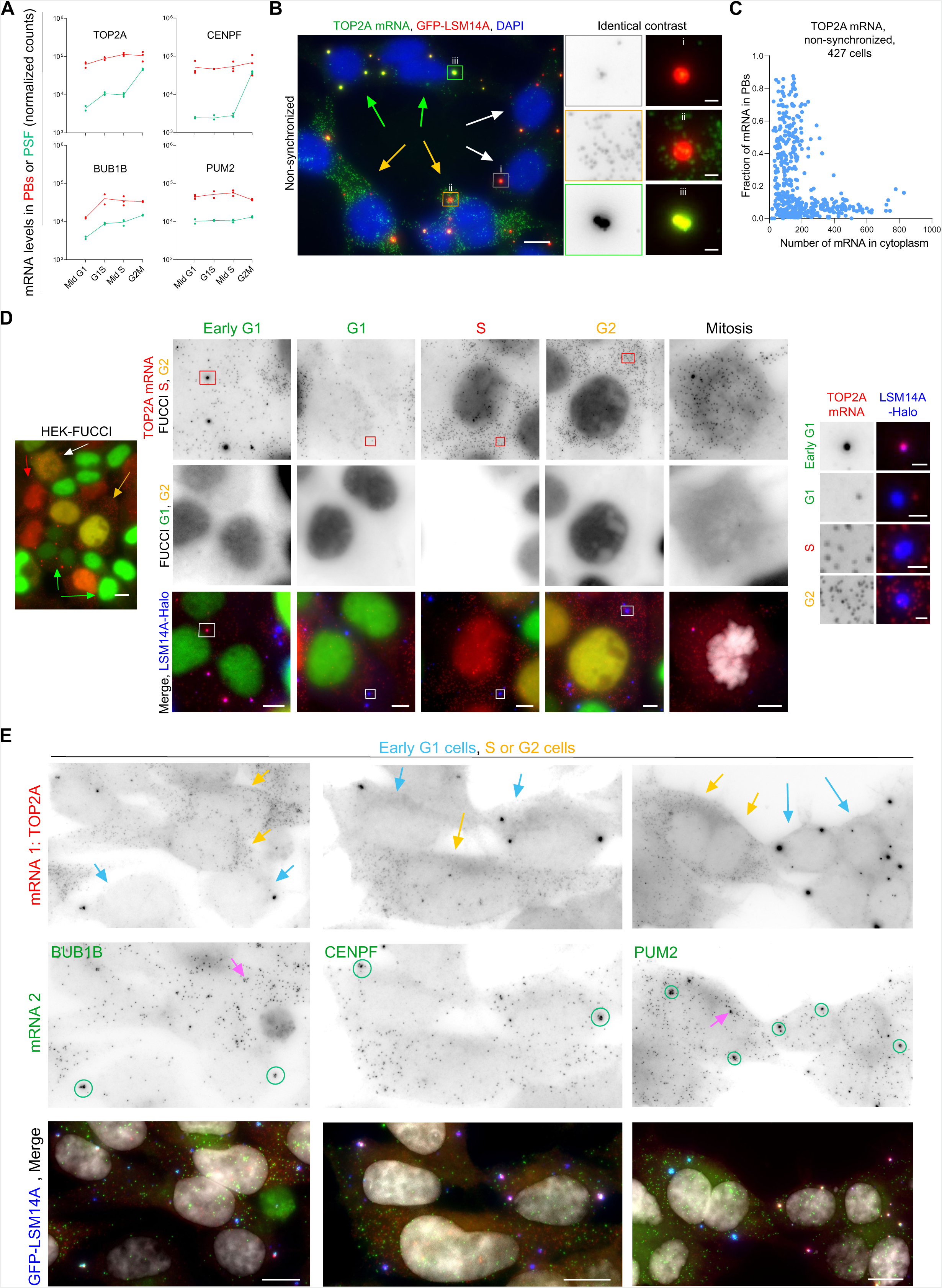
TOP2A, CENPF, BUB1B mRNAs concentrate in PBs after cell division. (A) Evolution of TOP2A, CENPF, BUB1B and PUM2 mRNA levels (in normalized counts) in purified PBs and pre-sorting fractions across the cell cycle. (B) Asynchronous HEK293-GFP-LSM14A cells after smFISH of TOP2A mRNA. The Cy3 smFISH signal is in green and GFP-LSM14A in red. Nuclei were stained with DAPI (blue). Insets present enlargements of representative PBs, illustrating the intercellular heterogeneity of RNA concentration in PBs. (C) Fraction of TOP2A mRNAs localized in PBs as a function of their number in the cytoplasm. Each dot corresponds to one cell. (D) Left panel: HEK-FUCCI cells transiently expressing LSM14A-Halo to label PBs (in far-red, shown in blue), after smFISH of TOP2A mRNA (Cy3, in red). The green, red, yellow, and white arrows point to cells in G1, S, G2, and M, respectively, identified based on the FUCCI system. These cells are enlarged in the middle panels. Insets on the right present enlargements of the representative PBs framed in the middle panels. (E) 2-color smFISH of TOP2A mRNAs and BUB1B, CENPF or PUM2 mRNAs in asynchronous HEK293-GFP-LSM14A cells. In the merge, TOP2A mRNAs (in Cy5, upper row) are in red, co-detected mRNAs (in Cy3, middle row) are in green, PBs are in blue, and DAPI-staining of the nuclei in white. Blue arrows point to early G1 cells and yellow ones to S or G2 cells, as revealed by the TOP2A mRNA labeling pattern. Green circles surround mRNA clustered in PBs. The pink arrows point to nuclear transcription sites. Scale bars: 10 µm in panels, 1 µm in insets.

Looking for mRNAs with similar expression patterns, we identified the mRNAs encoding the centromere protein F (CENPF) and, to a lesser extent, the mitotic checkpoint serine/threonine kinase B (BUB1B) (Figure 4A). Like TOP2A, these mRNAs are most highly expressed in G2 and encode proteins functioning mainly in G2 and mitosis^49–52^. Using 2-color smFISH, we found that CENPF and BUB1B mRNAs had similar dynamics as TOP2A: they co- localized with it in PBs in early G1 cells and were highly expressed yet weakly present in PBs in S or G2 cells (Figure 4E). As a benchmark, we used PUM2 mRNA, which displayed a stable expression and PB localization throughout the cell cycle (Figure 4A) and in the 2-color smFISH (Figure 4E). In summary, a group of transcripts encoding G2-M proteins traffic into PBs after cell division, when their protein is no longer needed, further highlighting the uncoupling between PB and cytoplasmic mRNA contents.

### A subset of mRNAs decaying at the mitosis-to-G1 transition accumulate in PBs in G1

Independently of the FAPS approach, we also explored RNA localization in PBs across the cell cycle using high-throughput smFISH (HT-smFISH, Figures 5A and S5). We selected 94 mRNAs with or without known cyclic expression and carried out their analysis in non-synchronized HEK293-GFP-LSM14A cells. Cells were classified in G1 or G2 phase based on their DAPI signal (see Methods, Figure 5A, Table S7). While most transcripts displayed a similar fraction in PBs in G1 and G2 cells, a subgroup of mRNAs showed higher accumulation in PBs in G1 than in G2 (Figure 5B). Their fraction in PBs was particularly heterogeneous in G1 cells, readily reaching values above 50% of the molecules (Figure 5C). Interestingly, expression of many of these transcripts was reported to peak in G2 and decrease in G1, while their encoded proteins are implicated in G2-to-mitosis progression (ECT2^53^, TTK^54^, and DLGAP5^55^).

**Figure 5:**
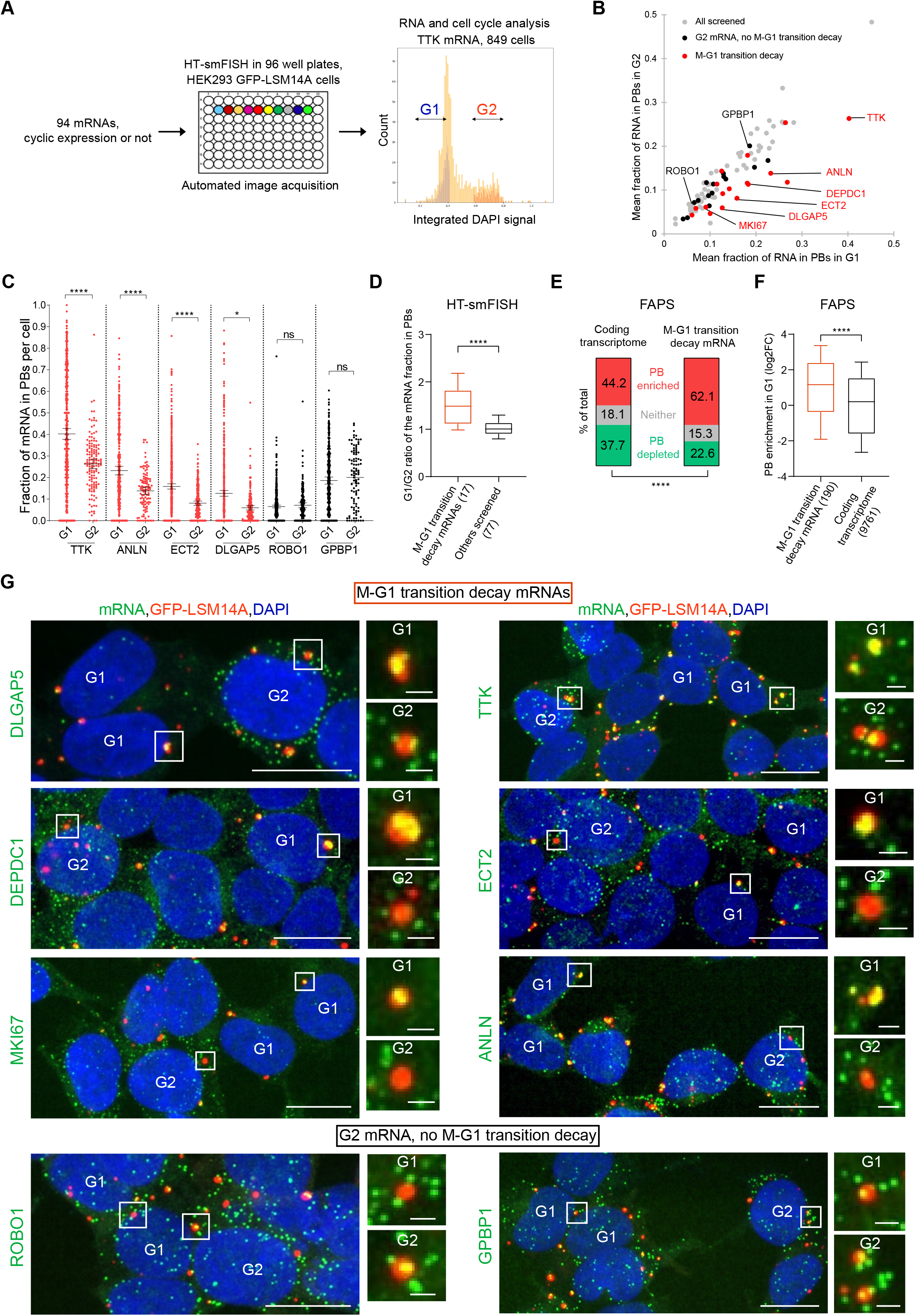
A high-throughput smFISH screen reveals PB localization of an mRNA subset after mitosis. (A) HT-smFISH workflow and typical cell cycle profiling based on DAPI staining. (B) Scatter plot comparing the fraction of mRNA molecules in PBs in G1 and G2 cells for the 94 screened mRNAs. The mRNAs decayed or not during M-G1 transition were taken from Krenning et al., 2022^22^. The indicated mRNAs are those shown in (G). (C) Fraction of mRNA molecules in PBs for indicated transcripts, in G1 and G2 cells. The number of analyzed cells is in Figure S5C. Horizontal lines, mean; error bars, 95% CI. Two-tailed Mann-Whitney test: ****, p<0.0001; *, p<0.05; ns, non-significant (p>0.05). (D) Boxplot showing the fold-change of the mRNA fraction in PBs in G1 versus G2 cells for mRNAs decaying at the M-G1 transition and other screened mRNAs. Two-tailed Mann-Whitney test: ****, p<0.0001. (E) Stacked bar graph showing the proportion of PB-enriched (red), PB-depleted (green), or neither (grey) mRNAs in G1 in the whole coding transcriptome and in M-G1 transition decay mRNAs (normalized counts >100). Chi-squared test: ****, p<0.0001. (F) Box plot showing PB enrichment in G1 for M-G1 transition decay mRNAs and for the whole coding transcriptome (normalized counts >100). Two-tailed Mann-Whitney test: ****, p<0.0001. (G) HEK293-GFP-LSM14A cells hybridized using HT-smFISH. Examples of mRNAs undergoing M-G1 transition decay or not. mRNA is shown in green, PBs in red, and nuclei in blue. DAPI-based cell cycle classification is indicated within cells. Scale bars: 10 µm in panels, 1 µm in insets.

We hypothesized that mRNA accumulation in PBs may be concomitant with their elimination after mitosis. In agreement, most mRNAs showing higher accumulation in PBs in G1 than in G2 belonged to a group of mRNAs previously reported to undergo decay at the mitosis-to-G1 (M-G1) transition^22^ (Figure 5B-D and G). Conversely, mRNAs encoding G2 proteins but not undergoing decay at the M-G1 transition accumulated similarly in PBs in G2 and G1 cells. To extend these data, we considered the 161 mRNAs described as decaying at the M-G1 transition^22^ and well-detected in our FAPS dataset. 62% of them were significantly enriched in PBs purified from G1 cells, which is higher than expected by chance (44%, Figure 5E). They also tended to reach higher levels of enrichment (Figure 5F). Thus, HT-smFISH independently provided evidence of cell cycle-dependent RNA localization in PBs, with mRNAs undergoing M- G1 transition decay particularly accumulating in PBs in G1.

### Cyclic PB localization does not result from fluctuating amounts of non-polysomal mRNAs

To explain cyclic mRNA localization in PBs, we first envisioned a mechanism related to the condensate nature of PBs. Indeed, it was previously proposed that condensates could attenuate cell-to-cell expression differences^56^. While the examples studied above clearly showed that mRNA localization in PBs does not limit expression changes in the cytoplasm throughout the cell cycle, we further challenged this hypothesis using the cell cycle-resolved TOP2A, FBXO5, and CLK1 smFISH data. Even within the same cell cycle phase, the fraction of mRNAs in PBs showed weak or no correlation with cytoplasmic expression (Figure S6A). This was true whether RNA fractions in PBs were high or low (e.g. TOP2A mRNAs in early G1 vs S or G2, respectively), and argued against a general role of PBs in recruiting excess cytoplasmic mRNAs.

Alternatively, PBs could simply recruit free mRNAs in the cytoplasm. In this scenario, the distinctive TOP2A, BUB1B and CENPF mRNA accumulation in PBs in early G1 could result from their specific translational downregulation after mitosis, and disrupting polysomes should be sufficient for their untimely recruitment in PBs in G2. However, treating HEK293-FUCCI cells transiently expressing LSM14A-Halo with puromycin, which disrupts polysomes within a few minutes^57^, did not cause TOP2A mRNAs to massively relocalize into PBs in G2 cells (Figure S6B-C). A 2-color smFISH of BUB1B or CENPF mRNAs with TOP2A mRNAs in HEK293-GFP- LSM14A cells showed a similar behavior (Figure 6A-D). In summary, disrupting polysomes was not sufficient to prematurely drive TOP2A, BUB1B, and CENPF mRNAs into PBs in G2.

**Figure 6:**
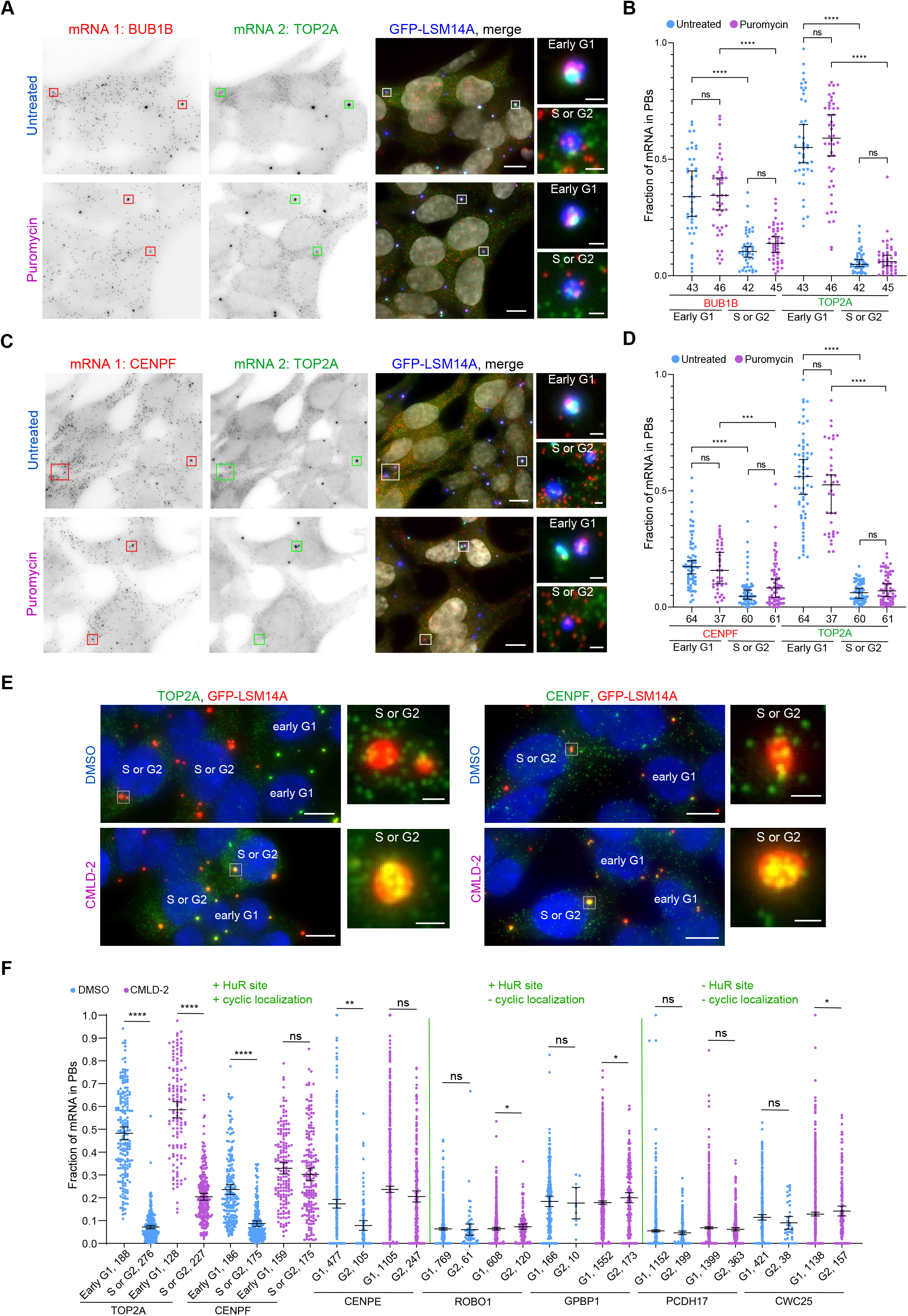
Determinants of cyclic localization of mRNAs in PBs. (A) 2-color smFISH of BUB1B and TOP2A mRNAs in asynchronous HEK293 cells expressing LSM14A-GFP, treated or not with puromycin for 1 hr. In the merge, BUB1B mRNAs (Cy3) are in red, TOP2A mRNAs (Cy5) in green, LSM14A-GFP-labelled PBs in blue and DAPI-staining of the nuclei in white. Cells were classified in early G1, or in S or G2 based on TOP2A mRNA labeling pattern. Scale bars: 10 µm in panels, 1 µm in insets. (B) Fraction of BUB1B and TOP2A mRNAs localized in PBs in early G1 and in S or G2, in control and puromycin-treated cells. Each dot corresponds to one cell (2 independent experiments). (C and D) Same as A and B for CENPF and TOP2A mRNAs. (E) Localization of TOP2A or CENPF mRNAs (in green) in HEK293-GFP-LSM14A cells after a 24h DMSO or CMLD-2 treatment. PBs are in red and nuclei in blue. (F) Fraction of test and control mRNAs localized in PBs in DMSO or CMLD-2-treated cells (2-3 independent experiments). TOP2A and CENPF were detected by smiFISH and the others by HT-smFISH. Horizontal lines, mean; error bars, 95% CI. Two-tailed Mann-Whitney tests: ****, p<0.0001; ***, p<0.0005; **, p<0.005; *, p<0.05; ns, non-significant (p>0.05). Scale bars: 10 µm in panels, 1 µm in insets.

### HuR-ARE interactions prevent mRNA localization in PBs in G2

We then searched for RBPs contributing to cyclic mRNA localization and noticed a particular overlap between 4E-T (a PB protein required for PB formation) and HuR (one of the ARE-binding proteins) targets. Analyzing the top 1000 PB-enriched mRNAs, we observed that targets of both 4E-T and HuR were less enriched in PBs in G2 than in the other phases, while targets of 4E-T but not HuR were more enriched in PBs in G2 (Figure S6D). A similar analysis of ARE-containing mRNAs showed this pattern was specific to HuR (Figure S6D). HuR shuttles between the nucleus and cytoplasm where it stabilizes cell cycle-related mRNAs^58,59^. Furthermore, its cytoplasmic fraction increases in S and G2 phases^59^, suggesting altogether that it could prevent mRNA localization in PBs in G2.

To test this, we relied on a chemical inhibitor (CMLD-2) which specifically disrupts HuR- ARE interactions^60^ and we imaged several G2 mRNAs reported as HuR targets and accumulating in PBs in G1 more than G2. Interestingly, a 24h treatment with CMLD-2 strikingly increased the fraction of mRNA localized in PBs in G2, abolishing (CENPE and CENPF) or reducing (TOP2A) the G2-G1 differential localization (Figure 6E, F). In contrast, control mRNAs, either not HuR targets or HuR targets without cyclic PB localization, were not affected by the drug (Figure 6E, F). In summary, we identified HuR as an RBP contributing to cyclic mRNA localization by preventing their recruitment in PBs during G2.

### Length favors accumulation of AU-rich mRNAs in PBs in early G1

We then searched for mRNA features that could participate in cyclic PB localization. We previously reported that in asynchronous cells, the GC content of CDS and 3’UTR is the best predictor of mRNA enrichment in PBs^8^. This held true at all cell cycle phases: PB-enriched mRNAs were particularly AU-rich, with a striking correlation between PB enrichment and GC content (r_s_ up to -0.79, Figure S7A, B). However, this compositional bias displayed only minor changes across the cell cycle, even when focusing on the top 1000 PB-enriched mRNAs (Figure S7C). Given the nucleotide bias in the CDS, NNA/U triplets were strongly overrepresented in PB-enriched mRNAs, as observed previously^8^ (Figure S7D). While similar to the codon usage bias reported for cell cycle-regulated genes^61^ (Figure S7E), it appeared unrelated to cell cycle-regulated PB localization (Figure S7D).

We also reported that PB-enriched mRNAs tend to be longer than average^8^. While this held true at all cell cycle phases (Figure S7F), the correlation between PB enrichment and mRNA length was higher in G1 than in the other phases (Figure S7G). The analysis of the top 1000 PB-enriched mRNAs further revealed that mRNAs were particularly long in mid G1 (Figure 7A). The size differences seemed to mainly originate from the CDS compared to 3’UTRs (Figure 7A). Similarly, mRNAs revealed by HT-smFISH to accumulate more in PBs in G1 than in G2, had particularly long CDS (Figure 7B). Strikingly, for the top 500 AU-rich mRNAs, mRNA length appeared as a key feature associated with PB localization in G1 (r_s_=0.55) but not in G2 (r_s_=- 0.12, Figure 7C).

**Figure 7:**
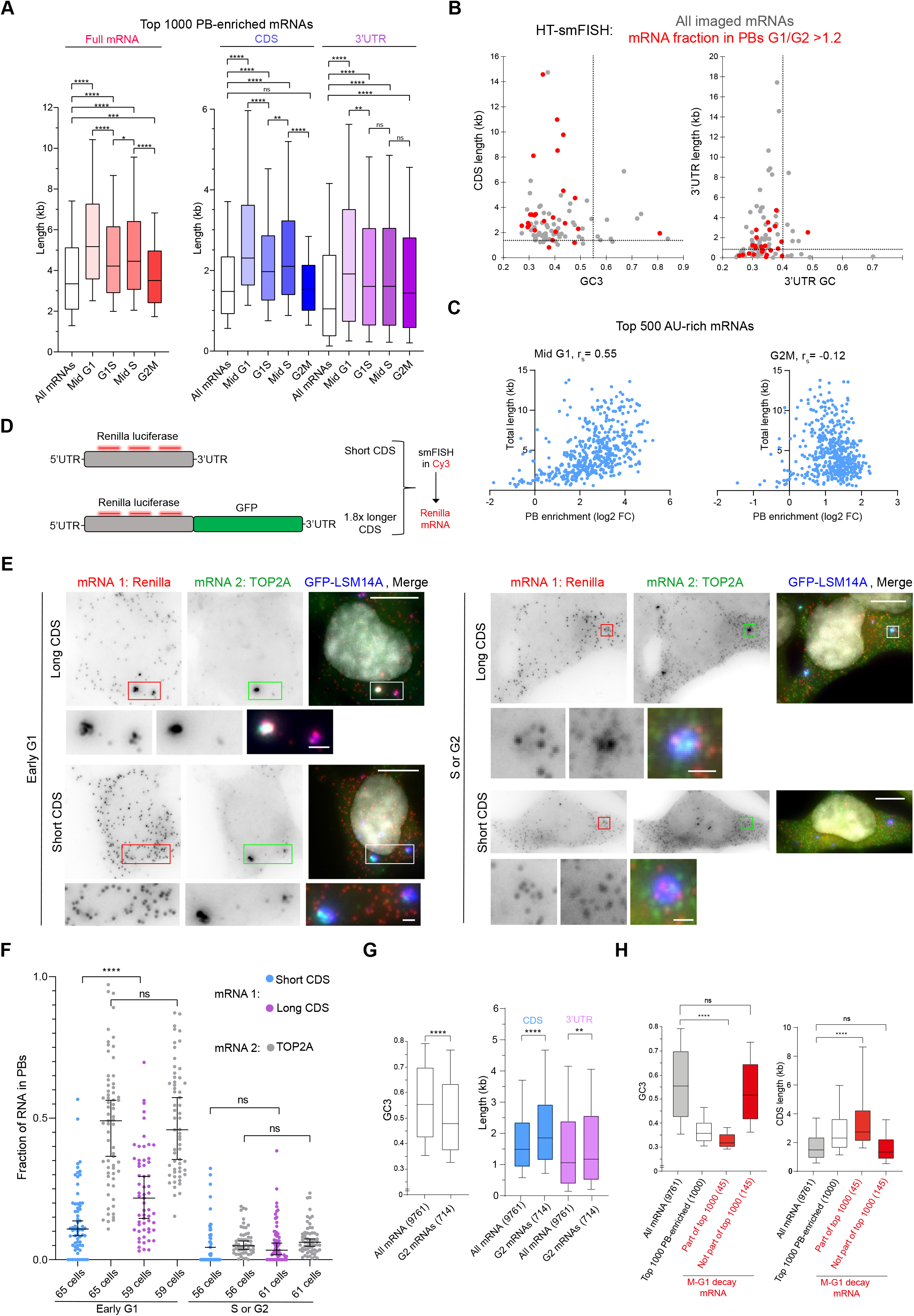
Length increases RNA accumulation in PBs in G1. (A) Lengths of the full mRNA, CDS, and 3’UTR for all mRNAs (normalized counts>100) or for the top 1000 PB-enriched mRNAs (FC>0, p-adj<0.05, PSF normalized counts>100) in the various cell cycle phases. Whiskers represent the 10 to 90% percentile. (B) CDS or 3’UTR length and GC content of the mRNAs imaged by HT-smFISH. mRNAs accumulating in PBs more in G1 than G2 are highlighted in red. Dotted lines indicate the median length and GC content of the coding transcriptome (PSF normalized counts>100). (C) Scatter plot of total length and PB enrichment of the top 500 AU-rich mRNAs, in mid G1 and G2M, with Spearman correlation (r_s_). (D) Schematic of the length reporter experiment, with smFISH probes in red. (E) 2-color smFISH of the reporter mRNA and TOP2A mRNA in asynchronous HEK293-GFP-LSM14A cells, after transfection of the long or short form of the Renilla Luciferase reporter. Renilla Luciferase mRNAs (Cy3) are in red, TOP2A mRNAs (Cy5) in green, GFP-LSM14A-labelled PBs in blue, and DAPI in white. Cells in early G1 and in S or G2 were classified based on TOP2A mRNA labeling pattern. Scale bars: 10 µm in panels, 1 µm in insets. (F) Fraction of long and short Renilla Luciferase mRNAs in PBs, with TOP2A mRNAs as an internal control (2 independent experiments). Horizontal lines, median; error bars, 95% CI. Two-tailed Mann-Whitney tests: ****, p<0.0001; **, p<0.005; *, p<0.05; ns, non-significant (p>0.05). (G and H) Features of G2M and M-G1 decay mRNAs compared to all mRNAs or the top 1000 PB-enriched mRNAs. Two-tailed Mann-Whitney tests: ****, p<0.0001; **, p<0.005; ns, non-significant (p>0.05).

We therefore transiently transfected Renilla Luciferase reporters of different length in HEK293-GFP-LSM14A cells and analyzed their localization using smFISH against Renilla along with TOP2A mRNAs to distinguish early G1 from S and G2 cells (Figure 7D, E). Importantly, all transcripts were similarly AU-rich (36-37% GC), making them amenable to PB localization^8^.

Lengthening the RLuc CDS from 0.9 to 1.7-1.8 kb using either GFP (Figure 7D-F) or RLuc itself (Figure S7H, I), led to significantly more accumulation in PBs in early G1 cells, while reporters minimally accumulated in PBs in S and G2 cells. The results were similar after introducing a stop codon between RLuc and GFP, which reversed the CDS length to 0.9 kb without affecting total length (Figure S7H, I). Therefore, the reporter assay pointed to the importance of total rather than CDS length for specific accumulation in PBs in early G1. This apparent discrepancy could be explained by the intrinsic features of G2-M mRNAs, among which many will accumulate in PBs in G1. First, they are particularly AU-rich (Figure 7G). Second, they are longer than average^28^, and we found it is due to their CDS more than their 3’UTR (Figure 7G).

Similarly, we analyzed the features of M-G1 decay mRNAs, which also accumulate in PBs in G1 (Figure 5). The M-G1 decay mRNAs that are part of the top1000 PB-enriched mRNAs in G1 tend to have long CDS and to be particularly AU-rich, like other mRNAs of this group (Figure 7H). In contrast, the M-G1 decay mRNAs not belonging to the top 1000-PB enriched mRNAs have average CDS length and nucleotide composition. Thus, long CDS and AU-rich composition are typical features of PB-enriched mRNAs in G1 rather than of M-G1 decay mRNAs. Taken together, these data show that mRNA length, in the context of an AU-rich nucleotide composition, is a feature contributing to RNA accumulation in PBs in G1 cells.

## Discussion

### The mRNA content of PBs is dynamic and partly uncoupled from the cytoplasmic content

In this study, we set out to investigate the dynamics of mRNA localization in PBs at the scale of hours in standard growth conditions. As a biological context, we chose the cell cycle since it entails changes in cytoplasmic RNA abundance that have been well documented at various RNA metabolism levels^13–15,18,19,21,22^. During every cell cycle, PBs form in G1, then increase in size more than in number, culminating in G2, before dissolving in mitosis. Cycling cells thus give access to the cell-autonomous dynamics of the PB content in the absence of any stress.

The genome-wide landscape of PB RNAs during cell cycle progression allowed answering two primary questions. Does the RNA content of PBs change during cell cycle progression? Do these changes simply mirror changes of cytoplasmic mRNA abundance, or do they behave distinctly? In fact, PBs undergo widespread changes in their mRNA content, and part of these changes did not reflect those of the cytoplasm: (i) mRNAs could change in PBs more strikingly than in the cytoplasm (particularly from mid G1 till mid S), (ii) and inversely, they could markedly change in the cytoplasm without a similar change in PBs (particularly from mid S to the next G1). Single molecule RNA imaging highlighted the remarkable variety in RNA localization patterns: for example, TOP2A, BUB1B and CENPF strikingly accumulated in PBs in early G1; many mRNAs peaking in G2 before undergoing decay accumulated in PBs in G1; CCNE2 at the G1S transition; FBXO5 in mid S; and CLK1 at the G2M transition. Thus, differential localization in PBs is not restricted to specific stages of the cell cycle. In particular, it does not follow the evolution of PBs considered as a whole (increasing size and number) during cell cycle progression.

### Biological relevance of differential mRNA accumulation in PBs

For most mRNAs, a minority of molecules accumulates in PBs, arguing against a causative role of PBs in regulating cyclic gene expression. Yet, some transcripts including TOP2A strongly accumulated in PBs in early G1. These transcripts are transcriptionally and translationally upregulated from late S to mitosis, periods when their encoded protein is needed^46,47,50,52^, and then subject to mitotic decay pathways^22^. TOP2A mRNA for instance is degraded by a CCR4- NOT-dependent mechanism^22^. However, there is no mechanism for ensuring translation inhibition of residual transcripts in G1^22,62^. We hypothesize that PB localization could contribute to the silencing of such transcripts in early G1. These examples were further corroborated by HT-smFISH analysis, which revealed a group of mRNAs that specifically accumulate in PBs in G1 and that are also subject to mitotic decay^22^. Yet, decay in G1 was not systematically associated with accumulation in PBs in G1. Regardless of how and where their decay takes place (not accessible through smFISH); our data suggest that PBs could compartmentalize mRNAs away from the translation machinery at specific cell cycle phases.

### PBs do not buffer excess non-polysomal mRNAs

To explain cyclic mRNA localization in PBs, we first envisioned a mechanism inspired by the physical nature of membraneless organelles. Since liquid-liquid phase separation (LLPS) implies the condensation of molecules that are over their saturation concentration, membraneless organelles have been proposed as a buffering mechanism to limit cell-to-cell variability of intracellular concentration^56^. While this was observed for minimal condensates (nanoclusters) forming upon massive mRNA storage in C. elegans arrested oocytes^63^, our smFISH data did not support such a hypothesis for PBs. Our results are rather consistent with the conclusion that multicomponent cellular organelles like PBs and stress granules are not governed by a fixed saturation concentration^64^.

Then, since polysomal mRNAs are excluded from PBs^7^, we considered a scenario where mRNA accumulation in PBs is governed by the amount of non-translating mRNAs. From a transcriptome-wide perspective, it has been previously proposed that non-optimal codon usage could generate cell cycle-dependent translation efficiency^61^. In all cell cycle phases, we found that PB-enriched mRNAs were particularly AU-rich, leading to a non-optimal codon usage associated with inefficient translation, as described previously^8^. However, neither their GC content, nor their codon usage differed between phases, indicating that these features are not involved in the cyclic pattern of mRNA accumulation in PBs. We also addressed the question experimentally. Releasing TOP2A, BUB1B and CENPF mRNAs from polysomes in G2 cells was not sufficient to relocate them in PBs, despite them having appropriate intrinsic features to accumulate in PBs in early G1. Altogether, we did not find evidence that mRNA accumulation in PBs represents the inverted mirror of mRNA translation.

### Intrinsic and extrinsic factors regulate cyclic localization in PBs

This prompted us to look for other determinants of cyclic mRNA localization in PBs. In terms of RBPs, we found that inhibiting HuR-ARE interaction causes the premature localization of HuR target mRNAs to PBs in S or G2. This highlights a role for HuR in preventing mRNA recruitment in PBs in later stages of the cell cycle. In terms of mRNA features, we previously reported that mRNA accumulation in PBs in asynchronous cells moderately correlates with their length^8^. To our surprise, this feature was cell cycle-dependent, with PB mRNAs longer in G1 than in G2.

Moreover, for the most AU-rich mRNAs of the transcriptome, PB enrichment correlated with length in G1 but not G2. Such a phase specific impact of mRNA length was confirmed using mRNA reporters. Overall, two mRNA features appear associated with PB localization: the strongest is their GC content, in all phases; the second is their length, specifically in G1.

Interestingly, *in-silico* modeling showed that longer transcripts localize at the core of RNA-protein condensates, where they augment the density of molecular interactions. This promotes and stabilizes condensates, preventing them to collapse due to excessive RNA recruitment^65,66^. It is particularly tempting to speculate that long RNAs favor the seeding and stabilize nascent PBs in early G1. These PBs could then progressively grow into ’’mature’’ PBs without collapsing. Long RNAs may at a certain point lose their advantage, because the interaction network becomes saturated, or also because the short RNAs at the surface^65^ provide a barrier to long RNA recruitment^67^.

Taken together, mRNA accumulation in PBs appears to be a multifaceted process involving the cumulative effect of extrinsic (HuR) and intrinsic (GC content, length) factors. If PBs are dynamic in a cell-autonomous process like cell cycle progression, they could be even more so across a whole organism, or during development and differentiation.

### Limitations of the study

For feasibility reasons, PBs were purified following cell cycle synchronization procedures, which could entail changes in gene expression. However, the majority of in situ RNA imaging experiments was performed on non-synchronized cells. Second, it wasn’t possible to perform deep and quantitative proteomic analysis from purified PBs, due to insufficient material. While our gross mass spectrometry analysis did not reveal drastic loss of the main known PB proteins at particular cell cycle phases, the observed variations of their mRNA content strongly suggest associated changes in terms of RBP content. Third, puromycin being a global inhibitor of translation elongation, may have a different impact on PB recruitment compared to endogenous transcript-specific translation initiation repression. Finally, further studies will be required to identify other factors that add to HuR binding, mRNA nucleotide composition and length, to control cell cycle-dependent localization in PBs.

## Acknowledgments

A.S., E.C. and A.L. were supported by the Agence Nationale pour la Recherche (ANR) grant number ANR-19-CE12-0024-01. The work was supported by grants from the Association pour la Recherche sur le Cancer (PJA20181208011) to D.W., ANR-19- CE12-0024 to D.W., D.G. and E.B., INCa grant 2022-082 to D.W., MSDAvenir to E.B., ANR-19-P3IA-0001 (PRAIRIE 3IA Institute) to T.W., and by the France Génomique national infrastructure, funded as part of the "Investissements d’Avenir" program managed by the ANR (contract ANR- 10-INBS-0009). We thank Nancy Standart (University of Cambridge, UK) for critical reading of the manuscript, Zoher Gueroui (École normale supérieure Paris, France) for scientific discussions, and Vincent Galy (IBPS Paris, France) for maintenance of the Zeiss microscope

## Author contributions

Conceptualization, A.S. and D.W.; Formal Analysis, A.S., M-N.B., F.S., M.K., A.L. and D.W.; Funding Acquisition, D.W.; Investigation, A.S., M-N.B., M.E.-L., E.C., A.M.G., A.V., C.B. and G.C.; Project Administration, D.W.; Software, T.B. and A.I.; Supervision, A.S., E.B., D.G., T.W., M.B. and D.W.; Validation, A.S., M-N.B., O.P., M.E.-L. and D.W.; Visualization, A.S., M-N.B. and M.K.; Writing – Original Draft, A.S. and D.W. All authors discussed the results and commented on the manuscript.

## STAR Methods

### Resource availability

#### Lead contact

Further information and requests for resources and reagents should be directed to and will be fulfilled by the lead contact, Dominique Weil

### Materials availability

All unique reagents generated by this study are available from the lead contact either without restriction or with a completed Materials Transfer Agreement.

### Data and code availability

- Raw RNA-seq data is available at ArrayExpress under the accession number E-MTAB- 12923
- All original code has been deposited at Zenodo and is publicly available as of the date of publication. DOIs are listed in the key resources table. The code used to analyze RNA PB localization is available at DOI: 10.5281/zenodo.12742387 or https://github.com/15bonte/p_bodies_cycle_2023 for standard smFISH experiments, and at DOI: 10.5281/zenodo.12666004 or https://github.com/Flo3333/Cell-cycle-and-HT-smFISH-analysis-of-RNA-localization-in-PBs for HT-smFISH analysis.
- Raw microscopy images and blots for all figures were deposited on Mendeley Data at DOI:10.17632/67s7c3dyc9.1
- Any additional information required to reanalyze the data reported in this paper is available from the lead contact upon request.

### Experimental model and subject details

#### Cell culture

HEK293 and HeLa cells were grown in Dulbecco’s modified Eagle’s Medium (DMEM, Gibco) supplemented with 10% fetal bovine serum (FBS, Sigma-Aldrich), and 100 U/mL penicillin/streptomycin (Sigma-Aldrich). HEK293-GFP-LSM14A cells were obtained by transfecting HEK293 cells with a pCMV-GFP-LSM1A plasmid and selecting stable clone under 500 μg/mL G418 (Gibco). HEK293-FUCCI cells were obtained via CRISPR-Cas9-mediated insertion of the PIP-FUCCI reporter in the AAVS1 genomic locus and selection under 1 μg/mL puromycin (Sigma-Aldrich). All cells were grown at 37 °C with 5% CO2.

### Method details

#### Cell cycle determination, drug treatments, and transfections

PIP-FUCCI allows accurate determination of cell cycle phases by expressing decay-sensitive fragments of two known cyclic proteins fused to fluorescent markers: (i) Cdt_11–17_- mVenus expressed in G1, G2 and mitosis (G2/M); and (ii) mCherry-Gem_1-110_ expressed in S and G2/M phases^68,69^. Both markers are fused with nuclear localization elements to make them compatible with cytoplasmic co-labeling. The accurate cell cycle-dependent expression of PIP-FUCCI was verified by flow cytometry analysis of live cells after DNA labeling with Hoechst. The mVenus positive, mCherry positive, and double positive cells showed the expected quantity of DNA (1N to 2N) for cells in G1, S, and G2/M phases, respectively (Figure S1A, B).

To inhibit translation, cells were treated with 100 μg/mL puromycin for 1 hr or 15 min, as indicated. The efficiency of the puromycin batch was controlled using HeLa cells expressing endogenous ASPM-MS2 mRNA that localizes to centrosomes in a translation-dependent manner^70,71^. As expected, centrosomal localization of this mRNA was abolished after a 15 min puromycin treatment (Figure S6C). To inhibit HuR-ARE interactions, cells were treated with 20 μM CMLD-2 (MedChemExpress) for 24 h. Control cells were treated with DMSO (Sigma-Aldrich) for the same time.

For all transient transfections, 5 μl Lipofectamine 2000 (Thermo Fisher Scientific) was used with the following DNA quantities per well of a 6-well plate: 1 μg LSM14A-Halo tag, and 100 ng of each Renilla luciferase construct with 900 ng of a non-transcribed plasmid. All transfection were done for 5 hr after which the reagent was washed off and cells allowed to grow for 24-48 hr before fixation. Cell fixation was done with 4% paraformaldehyde PFA (Electron Microscopy Sciences) for 20 min at room temperature (25°C). The Renilla Luciferase plasmids were expressed under a CMV promoter.

### Synchronizations

All synchronizations were performed in HEK293-GFP-LSM14A cells. For PB purification, synchronizations were done in 15 cm dishes. For double thymidine blocks, 4 million cells were seeded. The next day, 7.5 mM thymidine was added to the culture medium for 18 hr. Cells were then washed gently with warm PBS three times and further incubated in fresh complete medium. Cells were released for 9 hr after which 7.5 mM thymidine was added again for another 18 hr. This results in cells blocked at the G1S transition. To obtain cells in mid S phase, cells were further washed three times in warm PBS after which fresh complete medium was added. Cells were released for 5 hr to obtain a population at mid S phase. For CDK1 inhibition, 8 million cells were seeded. The next day the CDK1 inhibitor RO-3306 was added at 9 μM. After 22 hr, cells blocked at the G2M transition were harvested. To obtain cells at mid G1, we found that half the drug’s concentration (with the same blockage time) provides a better release from G2M into mid G1 (with three PBS washes and an 8 hr release in fresh complete medium). Forty to fifty 15 cm dishes for each cell cycle phase were synchronized. After each synchronization, a small fraction of each 15 cm dish was tested in flow cytometry for the quality of the synchronization and, if synchronization reached 55% for mid G1, 80% for G1S or mid S, and 70% for G2M, the rest was used for PB purification. For microscopy experiments, identical synchronizations were done for cells grown on 22x22 cm coverslips in 6-well plates, with the addition of poly-L-lysine for better cell adhesion.

For RNA-seq, 6 independent synchronization experiments were performed to obtain a sufficient number of cells in mid G1, G1S, G2M, and 7 for mid S. Each synchronization experiment was performed on 6-8 15 cm cell plates. For each FAPS, at least 2 synchronization experiments were pooled together. For each cell cycle stage, 2-3 days FAPS were required to purify enough material for three independent RNA extractions.

### Cell cycle related flow cytometry analysis and FACS purification

To fix cells for flow cytometry analysis, ∼100,000 cells were washed twice in PBS, trypsinized, and suspended in PBS. Cells were pelleted at 500 g for 5 min at 4°C and the pellet resuspended in 300 μl ice-cold PBS to which 900 μl cold ethanol (100% vol/vol) was slowly added. Cells were stored at -20°C overnight to allow fixation. The next day, cells were pelleted at 500 g for 5 min at 4°C. To label DNA, cells were resuspended in a 500 μl solution of 20 μg/mL RNaseA and 50 μg/mL propidium iodide in PBS and incubated at 37°C for 30 min. Flow cytometry was performed on a MACSQuant® Analyzer 10 Flow Cytometer (Miltenyi Biotec) using the 561 nm laser and acquisitions analyzed using FlowJo (v10.6.2). For flow cytometry analysis of living HEK293-FUCCI cells: cells grown to 80% confluency in 6-well plates were washed three times in PBS, trypsinized, and suspended in PBS. Cells were then pelleted for 5 min at 500 g at room temperature (25°C). The pellet was then resuspended in complete DMEM medium containing 15 µg/mL Hoechst 33342 (Life technologies) and incubated for 30 min at 37°C before flow cytometry analysis. Cytometry and analysis were performed using the same hardware (with a 405 nm laser) and software.

To sort HEK293-FUCCI cells by FACS: cell were grown to 80% confluency in 10 cm dishes and processed identically, minus the Hoechst 33342 staining. Around 1.5-2 million cells were sorted for each cell cycle phase (G1, S, or G2/M) using a MoFlo Astrios EQ (Beckman-Coulter, 488 and 561 nm lasers) and collected cells were pelleted for 5 min at 500 g. Cell pellets were flash frozen every 30 min to minimize the amount of cells transitioning into the next cell cycle phase and accumulated for western blotting.

### PB purification by FAPS

PB purification by FAPS is based on particle size and PB fluorescence. It was performed as described previously^7^, with a few modifications. HEK293-GFP-LSM14A cells (synchronized or not) were allowed to grow to ∼70-80% confluency in 15 cm dishes. After two PBS washes, cells were scrapped off, collected in 2 ml Eppendorf tubes and pelleted at 500 g for 5 min. Pellets were flash frozen in liquid nitrogen and stored at -80°C until the day before sorting. Each cell pellet was then resuspended in 1.5 mL ice-cold lysis buffer containing 50 mM Tris pH 8, 1 mM EDTA, 150 mM NaCl, and 0.2% Triton X-100 supplemented with 65 U/mL RNase out (Life Technologies) and 2x protease inhibitor cocktail (Roche), pipetted a few times and kept on ice for 20 min. Then, nuclei were pelleted by centrifugation for 5 min at 500 g, and the supernatant containing cytoplasmic organelles, including PBs, was transferred to a new tube. After supplementation with 10 mM MgSO_4_, 1 mM CaCl_2_, and 4 U/mL RQ1 DNase (Promega), samples were incubated at room temperature (25°C) for 30 min while avoiding direct sources of light to avoid GFP fluorescence bleaching. Next, the samples were centrifuged at 10,000 g for 7 min at 4°C. The supernatant was gently aspirated and the pellet was resuspended in 30 μl lysis buffer supplemented with 40 U RNase out. This was called the pre-sorting fraction. A small aliquot (1-3 μL) was stained with ethidium bromide and visualized using widefield fluorescence microscopy for a reference image before sorting. For each cell cycle phase, 3 independent pre-sorting fractions were kept aside for RNA-seq and mass spectrometry analyses to compare with their sorted counterparts.

Fluorescence activated particle sorting (FAPS) was carried out on a MoFlo Astrios EQ (Beckman-Coulter) using the 488 nm excitation laser. The PB sorting gate was determined using control samples of lysis buffer alone, or an identical cell preparation but made with cells expressing a truncated GFP-LSM14A protein that only displays diffuse fluorescence (GFP- LSM14A-Δ, described in ^7^) (Figure S1J). The pre-sorting sample was diluted 30 times in lysis buffer at the time of sorting (10 μL in 300 μL lysis buffer). The differential pressure during sorting was maintained around 0.8 and the sorting speed was around 8,000 events per second on average. The sorting purity was >95%. After sorting, a small aliquot was stained with ethidium bromide and visualized under widefield fluorescence microscopy. Purified PBs were pelleted at 10,000 g for 7 min in 2 mL low binding Eppendorf tubes, while the remaining was stored at - 80°C. ∼6 hr of active sorting were done per day and 2-3 days of sorting output were combined together to constitute a single replicate for RNA-seq for each cell cycle phase. A total of >140 hr of FAPS was needed to accumulate sufficient material for RNA-seq in triplicates and 1 mass spectrometry analysis for each of the 4 cell cycle conditions. Protein and RNA were extracted using TRIzol (Thermo Fisher Scientific). A total of ∼400 ng protein per condition and 5-10 ng RNA per replicate per condition were obtained.

### Mass spectrometry

Liquid chromatography–tandem mass spectrometry (LC-MS/MS) was performed at the proteomics platform at Institut Jacques Monod, Paris France. Proteins from the purified PB or pre-sorting fractions were dissolved in a 8 M urea solution (Sigma-Aldrich) before trypsin digestion. Samples were analyzed using an Orbitrap Fusion Lumos Tribrid Mass Spectrometer (Thermo Fisher Scientific) with the following settings: Ion Source: ESI (nano-spray), Fragmentation Mode: high energy CID, MS Scan Mode: FT-ICR/Orbitrap. Peptide and protein signals were processed using PEAKS Studio (v10.6 build 20201015) with the following parameters: Max Missed Cleavages: 2, Database: Homo Sapiens SwissProt Release_2020_06, Parent Mass Error Tolerance: 15.0 ppm, and Fragment Mass Error Tolerance: 0.5 Da. Protein and peptide signals were selected using a <1% false discovery rate filter. To classify proteins as PB-enriched or –depleted proteins, the Fisher’s exact test was performed in R. The significance cutoff was set to <0.025. The functional annotation used in Figures 1 and S2 was performed manually and is provided in Table S1.

### cDNA library generation and RNA sequencing

Library preparation and sequencing were performed at Ecole normale supérieure Génomique ENS core facility (Paris, France). cDNAs were synthesized using a combination of random primers and oligodT to amplify RNAs independently of their polyadenylation status. 2 ng of total RNA were amplified and converted to cDNA using Ovation RNA-Seq system v2 (TECAN). Following amplification, libraries were generated using the Nextera XT DNA Library Preparation Kit from Illumina. Libraries were multiplexed, and after single-end 75 bp sequencing (NextSeq 500, Illumina), 40 to 60 million reads per sample passed the Illumina filters. Three replicates per fraction and per cell cycle phase were made, for a total of 24 libraries.

### RNA-seq analysis

RNA-seq analysis was performed on a local Galaxy server. RNA STAR^72,73^ (version 2.7.8a) was used to align reads to the GRCh38p.13 assembly of the human genome with haplotypic duplications removed. The following parameters were used: 2 maximum mismatches per read alignment, a maximum ratio of mismatches to mapped length of 0.3, a minimum alignment score, normalized to read length of 0.66. All other parameters were set to default. More than 95% of reads mapped to the human genome, with the rest too short to be mapped. FeatureCounts^74^ (version 2.0.1) was used to count read using the Gencode v38 gene annotation. All parameters were set to default. Differential expression analysis was done using DEseq2 with the default settings. Normalized counts showed high reproducibility between replicates in both the PB and pre-sorting fractions across the cell cycle (Figure S4A), after exclusion of one PB replicate in G1S and one pre-sorting replicate in mid G1, due to lower reproducibility (R<0.9). Depending on the analysis, two normalizations were performed. (i) For PB enrichment calculation, PB replicates were normalized to pre-sorting replicates within each cell cycle phase, as performed previously^7^; (ii) to follow the evolution of RNA content in the PB and pre-sorting fractions across the cell cycle, all PB replicates were normalized together on the one hand, and all pre-sorting replicates together on the other hand. It can be noted that PB enlargement throughout interphase may lead to some underestimation of RNA content in PBs, since large PBs likely accumulate a larger pool of RNAs, while RNA-seq only measures relative RNA abundance.

### Gene ontology analysis

GO term analyses were performed using clusterProfiler^75,76^ in R. For GO analysis of PB mRNAs (Figure 2E), the reference list was a compilation of all mRNAs detected (normalized counts>0) in the PB fraction of at least one cell cycle phase, and the test lists contained mRNAs displaying differential abundance in the PB fraction (p-adj<0.05, DEseq2) between 2 cell cycle phases. Similarly, for GO analysis of the pre-sorted fraction mRNAs (Figure S3B, C), the reference list was a compilation of all mRNAs detected (normalized counts>0) in this fraction and the test lists contained the mRNAs differentially expressed (p-adj<0.05, DEseq2). The Benjamini & Hochberg p-value adjustment method was applied and the significance cutoff was set to <0.05. To limit GO term redundancy, simplified GO terms were used with a cutoff of 0.6 using the adjusted p-value, in Figure 2E. Dot plots were used to represent the top 10 GO categories with associated mRNAs count and p-adj values. Full detailed GO term lists are provided in Table S5.

### Immunofluorescence

Cells grown on 22x22 mm glass coverslips were fixed for 20 min at room temperature (25°C) with 4% PFA (Electron Microscopy Sciences) diluted in PBS and stored at 4°C. Before IF labeling, cells were permeabilized with 0.1% Triton X-100 in PBS (Sigma-Aldrich) for 15 min at room temperature (25°C). All primary and secondary antibodies were diluted in PBS 0.1% BSA. To label DDX6, we used a rabbit polyclonal anti-DDX6 antibody recognizing its C-ter extremity (BIOTECHNE, NB200-192, dilution 1/1000) and a secondary F(ab)2 goat anti-rabbit antibody labeled with AF488 (Life technologies, A11070, dilution 1/1000). To label the stress granule marker TIA1, we used goat polyclonal anti-TIA1 antibody (Santa Cruz, SC1751, dilution 1/200) and a secondary donkey anti-goat antibody labeled with Cy3 (Jackson ImmunoResearch, 705- 166-147, dilution 1/800). To label FXR1, we used rabbit polyclonal anti-FXR1 antibody (Sigma-Aldrich, HPA018246, dilution 1/500) and a secondary goat anti rabbit antibody labeled with AF488 (Life technologies, A11070, dilution 1/1000). To label FXR2, we used mouse monoclonal anti-FXR2 antibody (Life Technologies, MA1-16767, dilution 1/200) and a secondary goat anti mouse antibody labeled with AF488 (Life technologies, A11029, dilution 1/1000). Primary and secondary antibodies were incubated for 1 hr at room temperature after which coverslips were washed 3 times in PBS. Coverslips were then mounted using DAPI-containing Vectashield (Vector Laboratories).

### Single molecule FISH

Cells grown on 22x22 mm glass coverslips were fixed for 20 min at room temperature with 4% PFA (Electron Microscopy Sciences) diluted in PBS and permeabilized with 70% ethanol overnight at 4°C. We used the single molecule inexpensive variant of smFISH (smiFISH)^31^, which uses 24 primary probes for each RNA target each made of a gene-specific sequence that hybridizes to the RNA, and a common overhang that serve as platforms for recruiting fluorescently labeled oligos (either Cy3 or Cy5). To this end, 40 pmoles of primary probes were first pre-hybridized to 50 pmoles of fluorescently labeled oligos in 10 μL of 100 mM NaCl, 50 mM Tris-HCl, 10 mM MgCl_2_, pH 7.9. This was performed on a thermocycler with the following program: 85 °C for 3 min, 65 °C for 3 min, and 25 °C for 5 min. The resulting fluorescent probe duplexes were then used in a hybridization mixture consisting of: 1x SSC, 0.34 mg/mL tRNA, 15% formamide (Sigma-Aldrich), 2 mM vanadyl ribonucleoside complexes (Sigma-Aldrich), 0.2 mg/mL RNase-free bovine serum albumin (Roche Diagnostics), 10% Dextran sulfate (Eurobio), and 2 μL fluorescent duplexes per 100 μl hybridization volume. Hybridization was performed overnight at 37°C. The next day, coverslips were washed in a 15% formamide 1x SSC solution for 40 min twice. Coverslips were then mounted using DAPI-containing Vectashield (Vector Laboratories, Inc.). All probe sequence are available in Table S6.

### High-throughput smFISH

HT-smFISH was performed as described previously^77^. Probesets against mRNAs of interest were generated starting from a pool of DNA oligonucleotides (GenScript). The design of DNA oligonucleotides was based on the Oligostan script^31^. Briefly, oligos belonging to the same probeset (hybridizing to the same mRNA target) share a common barcode which allows their specific amplification through 2 rounds of PCR. Then, *in-vitro* transcription was used to generate transcript-specific primary RNA probesets. Each primary RNA probe contains a hybridization sequence recognizing the target of interest, flanked by 2 readout sequences. These readout sequences serve as platforms to recruit fluorescent TYE-563-labeled locked nuclei acids (similar to the smiFISH technique^31^). For each mRNA target, 25 ng of each of the fluorescent locked nuclei acids were prehybridized with 50 ng of primary RNA probeset in 100 μL of a solution containing 7.5 M urea (Sigma-Aldrich), 0.34 mg/mL tRNA, and 10% Dextran sulfate. Pre-hybridization was done using thermocycler with the following program: 90°C for 3 min, 53°C for 15 min, giving rise to fluorescent duplexes. Cells were grown in 96-well plates with glass bottoms, fixed with 4% PFA for 20 min, and permeabilized with 70% ethanol overnight. Cells were then washed with PBS and hybridization buffer (1× SSC, 7.5 M urea), and the 100 μL solution containing fluorescent duplexes was added. Cells were incubated at 48 °C overnight. The next morning, plates were washed 8 times for 20 min each in 1× SSC, and 7.5 M urea at 48 °C. Finally, cells were washed with PBs, labeled with DAPI at 5 μg/mL, and mounted in 90% glycerol (VWR), 1 mg/mL p-Phenylenediamine (Sigma-Aldrich) and PBS pH 8.

### Imaging

smFISH and IF imaging were performed using an inverted Zeiss Z1 widefield fluorescence microscope equipped with a motorized stage. A 63x oil objective with a numerical aperture of 1.4 and an Axiocam 506 mono camera (Zeiss) with a pixel size of 72 nm were used. The microscope was controlled by the Zeiss ZEN blue software (version 3.5.093.00003). Z stacks were acquired with a 0.3 μm spacing between each plane. This spacing provided adequate single molecule detection without oversampling. Maximum intensity projection (MIP) was used to obtain 2D images for visualization, mounting, and analysis. The following exposure duration and laser powers were used: 300-500 ms exposure at 100% laser power to image RNA in the Cy3 or Cy5 channels and PBs labeled with anti-DDX6 in the GFP channel; 50 ms at 50% laser power to image GFP-LSM14A-labeled PBs in the GFP channel and DNA in the DAPI channel. For the HT-smFISH screen, 96-well plates were imaged on an Opera Phenix High-Content Screening System (PerkinElmer), with a 63× water-immersion objective (NA 1.15). Three-dimensional images were acquired, with a spacing of 0.3 μm. Figures were mounted using Fiji^78^, Adobe illustrator, and the OMERO Figure tool^79^.

### Image analysis

To analyze PB characteristics in HEK293-FUCCI cells, nuclei and cell segmentation was done using the Cellpose^80^ model with watershed. Nuclei were segmented using the DAPI channel, and the cytoplasm using the DDX6 IF channel using an intensity threshold of 350-450. To determine the cell cycle phase based on the nuclear FUCCI signal, we used the Otsu method to classify nuclei as positive or negative in the green and red channels. All images were inspected to verify the reliability of the classification. PBs were detected based on DDX6 IF using the BigFISH^81^ Segmentation subpackage applied to the cytoplasm, and PB GFP masks were generated based on a fluorescence intensity threshold of 1200-1500. PBs were then assigned to individual cells and the mean PB fluorescence intensity was computed. The fluorescence background (250), estimated from areas of the image without cells, was subtracted from PB intensity values. PB size was computed as the maximal rectangular perimeter occupied by a PB, multiplied by the pixel size (72 nm) and divided by 2. PBs with size appearing smaller than 200 nm were excluded as they approach the diffraction limit of the microscope in the green channel. Images from 2 independent experiments were normalized to have the same PB intensity distribution means. The few PBs overlapping the nucleus in the MIPs were not taken into consideration.

smFISH images were analysed using BigFISH^81^, a python implementation of FISH- quant^82^ available at https://github.com/fish-quant/big-fish. The specific code used to analyze RNA localization in PBs is available at https://github.com/15bonte/p_bodies_cycle_2023. Nuclei and cell segmentation was done using the Cellpose^80^ model with watershed. Nuclei were segmented using the DAPI channel. The cytoplasm was segmented using background from the smFISH channel using an intensity threshold of 400-500. Spot detection was done using the Detection subpackage of BigFISH, which uses a Laplacian of Gaussian filter to enhance the spot signal. A local maximal algorithm then localizes every peak in the image, and a threshold was applied to discriminate the actual spots from the nonspecific background signal. One advantage BigFISH provides is the ability to automatically set an optimal RNA spot detection threshold regardless of the signal-to-background ratio of the smFISH. This parameter-free detection was used for all RNAs imaged in this study and manually adjusted when needed to obtain a unimodal distribution of single molecule fluorescence intensities. Next, we decomposed dense areas of RNA signal, firstly by removing background noise with a Gaussian background subtraction and secondly using the cluster decomposition tool. This tool computes the median detected spot intensity distribution, fits it with a Gaussian distribution signal, and uses it to compute the number of spots that best fit within each cluster. Next we detected PBs as masks using the GFP channel and the BigFISH Segmentation subpackage with a fluorescence intensity threshold of 2200 for cells synchronized in G1S, mid S and G2M, and 1600 for cells synchronized in mid G1, since PBs were less bright. We then computed two RNA populations: (i) RNA molecules in the cytoplasm (i.e. outside the nucleus), (ii) and RNA molecules in PBs (i.e. within the GFP mask). Since BigFISH was applied to MIPs, some PBs overlapped with the nucleus in a minority of cells and were excluded from the analysis. Cells without labeled PBs, with improper segmentation, or incomplete cell fragments were also excluded. In experiments involving transient transfection (Figure 7, S7, Renilla Luciferase reporters), cells with fewer than 15 mRNA molecules or with too high expression for proper BigFISH detection were excluded and the total mRNA was used to calculate the fraction of mRNA in PBs in cells with PBs appearing in the nucleus.

For HT-smFISH analysis (Figure 5, 6), a similar BigFISH pipeline was used to segment cells, nuclei and detect RNA spots. The specific code used to analyze HT-smFISH images is available here: https://github.com/Flo3333/Cell-cycle-and-HT-smFISH-analysis-of-RNA-localization-in-PBs. PBs were detected as 2D mask using a log filter (sigma = 2.25) to detect PB edges from the average projection of the image stack. A threshold was then applied to segment PBs edges by detecting positive gradients of intensity on the filtered image. Finally, PBs masks were filled and artifacts smaller than 10 pixels are removed. Cells without labeled PBs or RNA spot, with improper segmentation, and incomplete cells were excluded from the image analysis. To classify cells in G1 or G2, the DAPI signal was used to construct a cell cycle profile, as follows. We first calculated the integrated DAPI signal in each nucleus (nucleus area multiplied by the mean DAPI signal). The obtained profile was then fitted, using a ranking-based classification, to a cell cycle profile of HEK293-GFP-LSM14A cells obtained with flow cytometry and analyzed using the Dean-Jett model (∼50% cells in G1 and ∼16.5% of cells in G2, see example in Figure 5A).

### Western blots

Cytoplasmic proteins were extracted as described previously^83^, separated on a NuPage 4%– 12% gel (Invitrogen, Life Technologies) and transferred to Optitran BA-S83 nitrocellulose membrane (Fisher Scientific). After blocking in PBS containing 5% (wt/vol) nonfat dry milk for 1 hr at room temperature, membranes were incubated with the primary antibody overnight at 4°C, washed in PBS, and incubated with horseradish peroxidase–conjugated anti-rabbit secondary antibody (Interchim, Cat#111-036-003) diluted 1/10,000 for 1 hr at room temperature. Primary antibodies were: rabbit anti-DDX6 (BIOTECHNE, Cat# NB200-192, diluted 1/15,000), rabbit anti-4E-T (Abcam, Cat#ab95030, 1/2,500), rabbit anti-LSM14A (Bethyl, Cat#A305-102A, 1/5,000), rabbit anti-PAT1b (Cell Signaling, Cat#14288S, 1/1,000), and rabbit anti- rProt S6 (5G10, Cell signaling, Cat#2217, 1/5,000). After washing in PBS, proteins were detected using the Western lightning plus ECL kit (Perkin Elmer) and visualized by exposure to CL-XPosure film (Thermo Scientific).

### Statistical analysis

Graphical representations and statistical tests were performed using the GraphPad Prism software (v8, GraphPad software, Inc), the R suite (v 4.2.0, https://www.R-project.org, R Core Team 2018, R: A language and environment for statistical computing, R Foundation for Statistical Computing, Vienna, Austria), R studio (v 2022.12.0), and Excel 2016 and the Excel Analysis ToolPak (Microsoft). Venn diagrams were generated using a tool available at https://bioinformatics.psb.ugent.be/webtools/Venn. For imaging experiments, cell population distributions were compared using a Two-tailed Mann-Whitney test.

## Supplemental tables

**Table S1: Mass spectrometry analysis before and after sorting, related to Figure 1**. The proteomic data for each cell cycle phase are shown in independent tabs. The two other tabs provide the tentative list of PB proteins and the functional protein annotation used to build Figures 1 and S2.

Table S2: DEseq2 results comparing RNA-seq before and after sorting, related to Figure **2**. The normalized RNA expression levels (in normalized counts) for each replicate per cell cycle phase before and after sorting are shown, as well as their mean value.

**Table S3: DEseq2 results comparing RNA-seq before and after sorting throughout the cell cycle, related to Figure 2**. Every tab compares two successive cell cycle phases in the PB or the presorting fractions. The periodic phase of cyclic RNAs obtained from Dominguez et al., 2016 is also included^28^.

**Table S4: Normalized RNA expression levels (in normalized counts) before and after sorting across the cell cycle, related to Figure 2**. These correspond to the normalization used in Table S3. Values are shown for each replicate in both fractions throughout the cell cycle.

**Table S5: Gene ontology analysis results, related to Figure 2**. P-values, gene IDs, and gene counts are provided for each GO category shown in Figures 2E and S7B, C.

Table S6: Sequences of smFISH probes used throughout the study, related to Figures 3-7 and STAR Methods.

**Table S7: Data related to the HT-smFISH experiments, related to Figure 5**. RNA names, cell numbers, total mRNA counts, mRNA counts in PBs, mean fraction of mRNA in PBs, fraction of mRNA in PBs per cell, and decay status in G1 are indicated.

## References

1. Lyon, A.S., Peeples, W.B., and Rosen, M.K. (2021). A framework for understanding the functions of biomolecular condensates across scales. Nat Rev Mol Cell Biol 22, 215–235. 10.1038/s41580-020-00303-z.

2. Antifeeva, I.A., Fonin, A.V., Fefilova, A.S., Stepanenko, O.V., Povarova, O.I., Silonov, S.A., Kuznetsova, I.M., Uversky, V.N., and Turoverov, K.K. (2022). Liquid-liquid phase separation as an organizing principle of intracellular space: overview of the evolution of the cell compartmentalization concept. Cell Mol Life Sci 79, 251. 10.1007/s00018-022-04276-4.

3. Hirose, T., Ninomiya, K., Nakagawa, S., and Yamazaki, T. (2022). A guide to membraneless organelles and their various roles in gene regulation. Nat Rev Mol Cell Biol, 1–17. 10.1038/s41580-022-00558-8.

4. Woodruff, J.B., Hyman, A.A., and Boke, E. (2018). Organization and Function of Non-dynamic Biomolecular Condensates. Trends in Biochemical Sciences 43, 81–94. 10.1016/j.tibs.2017.11.005.

5. Bhat, P., Honson, D., and Guttman, M. (2021). Nuclear compartmentalization as a mechanism of quantitative control of gene expression. Nat Rev Mol Cell Biol 22, 653–670. 10.1038/s41580-021-00387-1.

6. Standart, N., and Weil, D. (2018). P-Bodies: Cytosolic Droplets for Coordinated mRNA Storage. Trends Genet. 34, 612–626. 10.1016/j.tig.2018.05.005.

7. Hubstenberger, A., Courel, M., Bénard, M., Souquere, S., Ernoult-Lange, M., Chouaib, R., Yi, Z., Morlot, J.-B., Munier, A., Fradet, M., et al. (2017). P-Body Purification Reveals the Condensation of Repressed mRNA Regulons. Mol. Cell 68, 144–157.e5. 10.1016/j.molcel.2017.09.003.

8. Courel, M., Clément, Y., Bossevain, C., Foretek, D., Vidal Cruchez, O., Yi, Z., Bénard, M., Benassy, M.- N., Kress, M., Vindry, C., et al. (2019). GC content shapes mRNA storage and decay in human cells. eLife 8, e49708. 10.7554/eLife.49708.

9. Pillai, R.S., Bhattacharyya, S.N., Artus, C.G., Zoller, T., Cougot, N., Basyuk, E., Bertrand, E., and Filipowicz, W. (2005). Inhibition of Translational Initiation by Let-7 MicroRNA in Human Cells. Science 309, 1573–1576. 10.1126/science.1115079.

10. Pitchiaya, S., Mourao, M.D.A., Jalihal, A.P., Xiao, L., Jiang, X., Chinnaiyan, A.M., Schnell, S., and Walter, N.G. (2019). Dynamic Recruitment of Single RNAs to Processing Bodies Depends on RNA Functionality. Molecular Cell 74, 521–533.e6. 10.1016/j.molcel.2019.03.001.

11. Bhattacharyya, S.N., Habermacher, R., Martine, U., Closs, E.I., and Filipowicz, W. (2006). Relief of microRNA-Mediated Translational Repression in Human Cells Subjected to Stress. Cell 125, 1111– 1124. 10.1016/j.cell.2006.04.031.

12. Aizer, A., Kalo, A., Kafri, P., Shraga, A., Ben-Yishay, R., Jacob, A., Kinor, N., and Shav-Tal, Y. (2014). Quantifying mRNA targeting to P-bodies in living human cells reveals their dual role in mRNA decay and storage. J Cell Sci 127, 4443–4456. 10.1242/jcs.152975.

13. Cho, C.-Y., Kelliher, C.M., and Haase, S.B. (2019). The cell-cycle transcriptional network generates and transmits a pulse of transcription once each cell cycle. Cell Cycle 18, 363–378. 10.1080/15384101.2019.1570655.

14. Sun, Q., Jiao, F., Lin, G., Yu, J., and Tang, M. (2019). The nonlinear dynamics and fluctuations of mRNA levels in cell cycle coupled transcription. PLOS Computational Biology 15, e1007017. 10.1371/journal.pcbi.1007017.

15. Bristow, S.L., Leman, A.R., and Haase, S.B. (2014). Cell Cycle-Regulated Transcription: Effectively Using a Genomics Toolbox. Cell Cycle Control 1170, 3–27. 10.1007/978-1-4939-0888-2_1.

16. Liu, Y., Chen, S., Wang, S., Soares, F., Fischer, M., Meng, F., Du, Z., Lin, C., Meyer, C., DeCaprio, J.A., et al. (2017). Transcriptional landscape of the human cell cycle. Proceedings of the National Academy of Sciences 114, 3473–3478. 10.1073/pnas.1617636114.

17. Fischer, M., Schade, A.E., Branigan, T.B., Müller, G.A., and DeCaprio, J.A. (2022). Coordinating gene expression during the cell cycle. Trends in Biochemical Sciences 47, 1009–1022. 10.1016/j.tibs.2022.06.007.

18. Bertoli, C., Skotheim, J.M., and de Bruin, R.A.M. (2013). Control of cell cycle transcription during G1 and S phases. Nat Rev Mol Cell Biol 14, 518–528. 10.1038/nrm3629.

19. Eser, P., Demel, C., Maier, K.C., Schwalb, B., Pirkl, N., Martin, D.E., Cramer, P., and Tresch, A. (2014). Periodic mRNA synthesis and degradation co-operate during cell cycle gene expression. Molecular Systems Biology 10, 717. 10.1002/msb.134886.

20. Chávez, S., García-Martínez, J., Delgado-Ramos, L., and Pérez-Ortín, J.E. (2016). The importance of controlling mRNA turnover during cell proliferation. Curr Genet 62, 701–710. 10.1007/s00294-016-0594-2.

21. Catala, M., and Abou Elela, S. (2019). Promoter-dependent nuclear RNA degradation ensures cell cycle-specific gene expression. Commun Biol 2, 1–13. 10.1038/s42003-019-0441-3.

22. Krenning, L., Sonneveld, S., and Tanenbaum, M.E. (2022). Time-resolved single-cell sequencing identifies multiple waves of mRNA decay during the mitosis-to-G1 phase transition. eLife 11, e71356. 10.7554/eLife.71356.

23. Battich, N., Beumer, J., de Barbanson, B., Krenning, L., Baron, C.S., Tanenbaum, M.E., Clevers, H., and van Oudenaarden, A. (2020). Sequencing metabolically labeled transcripts in single cells reveals mRNA turnover strategies. Science 367, 1151–1156. 10.1126/science.aax3072.

24. Gingold, H., Tehler, D., Christoffersen, N.R., Nielsen, M.M., Asmar, F., Kooistra, S.M., Christophersen, N.S., Christensen, L.L., Borre, M., Sørensen, K.D., et al. (2014). A dual program for translation regulation in cellular proliferation and differentiation. Cell 158, 1281–1292. 10.1016/j.cell.2014.08.011.

25. Yang, Z., Jakymiw, A., Wood, M.R., Eystathioy, T., Rubin, R.L., Fritzler, M.J., and Chan, E.K.L. (2004). GW182 is critical for the stability of GW bodies expressed during the cell cycle and cell proliferation. Journal of Cell Science 117, 5567–5578. 10.1242/jcs.01477.

26. Ayache, J., Bénard, M., Ernoult-Lange, M., Minshall, N., Standart, N., Kress, M., and Weil, D. (2015). P-body assembly requires DDX6 repression complexes rather than decay or Ataxin2/2L complexes. MBoC 26, 2579–2595. 10.1091/mbc.E15-03-0136.

27. Vassilev, L.T., Tovar, C., Chen, S., Knezevic, D., Zhao, X., Sun, H., Heimbrook, D.C., and Chen, L. (2006). Selective small-molecule inhibitor reveals critical mitotic functions of human CDK1. Proceedings of the National Academy of Sciences 103, 10660–10665. 10.1073/pnas.0600447103.

28. Dominguez, D., Tsai, Y.-H., Gomez, N., Jha, D.K., Davis, I., and Wang, Z. (2016). A high-resolution transcriptome map of cell cycle reveals novel connections between periodic genes and cancer. Cell Res 26, 946–962. 10.1038/cr.2016.84.

29. Love, M.I., Huber, W., and Anders, S. (2014). Moderated estimation of fold change and dispersion for RNA-seq data with DESeq2. Genome Biol 15, 1–21. 10.1186/s13059-014-0550-8.

30. Femino, A.M., Fay, F.S., Fogarty, K., and Singer, R.H. (1998). Visualization of Single RNA Transcripts in Situ. Science 280, 585–590. 10.1126/science.280.5363.585.

31. Tsanov, N., Samacoits, A., Chouaib, R., Traboulsi, A.-M., Gostan, T., Weber, C., Zimmer, C., Zibara, K., Walter, T., Peter, M., et al. (2016). smiFISH and FISH-quant - a flexible single RNA detection approach with super-resolution capability. Nucleic Acids Res. 44, e165. 10.1093/nar/gkw784.

32. Cappell, S.D., Mark, K.G., Garbett, D., Pack, L.R., Rape, M., and Meyer, T. (2018). EMI1 switches from being a substrate to an inhibitor of APC/CCDH1 to start the cell cycle. Nature 558, 313–317. 10.1038/s41586-018-0199-7.

33. Reimann, J.D., Gardner, B.E., Margottin-Goguet, F., and Jackson, P.K. (2001). Emi1 regulates the anaphase-promoting complex by a different mechanism than Mad2 proteins. Genes Dev 15, 3278– 3285. 10.1101/gad.945701.

34. Machida, Y.J., and Dutta, A. (2007). The APC/C inhibitor, Emi1, is essential for prevention of rereplication. Genes Dev 21, 184–194. 10.1101/gad.1495007.

35. Reimann, J.D., Freed, E., Hsu, J.Y., Kramer, E.R., Peters, J.M., and Jackson, P.K. (2001). Emi1 is a mitotic regulator that interacts with Cdc20 and inhibits the anaphase promoting complex. Cell 105, 645–655. 10.1016/s0092-8674(01)00361-0.

36. Miller, J.J., Summers, M.K., Hansen, D.V., Nachury, M.V., Lehman, N.L., Loktev, A., and Jackson, P.K. (2006). Emi1 stably binds and inhibits the anaphase-promoting complex/cyclosome as a pseudosubstrate inhibitor. Genes Dev 20, 2410–2420. 10.1101/gad.1454006.

37. Di Fiore, B., and Pines, J. (2007). Emi1 is needed to couple DNA replication with mitosis but does not regulate activation of the mitotic APC/C. Journal of Cell Biology 177, 425–437. 10.1083/jcb.200611166.

38. Hsu, J.Y., Reimann, J.D.R., Sørensen, C.S., Lukas, J., and Jackson, P.K. (2002). E2F-dependent accumulation of hEmi1 regulates S phase entry by inhibiting APC(Cdh1). Nat Cell Biol 4, 358–366. 10.1038/ncb785.

39. Dominguez, D., Tsai, Y.-H., Weatheritt, R., Wang, Y., Blencowe, B.J., and Wang, Z. (2016). An extensive program of periodic alternative splicing linked to cell cycle progression. eLife 5, e10288. 10.7554/eLife.10288.

40. Nielsen, C.F., Zhang, T., Barisic, M., Kalitsis, P., and Hudson, D.F. (2020). Topoisomerase IIα is essential for maintenance of mitotic chromosome structure. Proc Natl Acad Sci U S A 117, 12131– 12142. 10.1073/pnas.2001760117.

41. Lee, S., Jung, S.-R., Heo, K., Byl, J.A.W., Deweese, J.E., Osheroff, N., and Hohng, S. (2012). DNA cleavage and opening reactions of human topoisomerase IIα are regulated via Mg2+-mediated dynamic bending of gate-DNA. Proc Natl Acad Sci U S A 109, 2925–2930. 10.1073/pnas.1115704109.

42. Roca, J. (2009). Topoisomerase II: a fitted mechanism for the chromatin landscape. Nucleic Acids Res 37, 721–730. 10.1093/nar/gkn994.

43. Heck, M.M., Hittelman, W.N., and Earnshaw, W.C. (1988). Differential expression of DNA topoisomerases I and II during the eukaryotic cell cycle. Proc Natl Acad Sci U S A 85, 1086–1090. 10.1073/pnas.85.4.1086.

44. Woessner, R.D., Mattern, M.R., Mirabelli, C.K., Johnson, R.K., and Drake, F.H. (1991). Proliferation- and cell cycle-dependent differences in expression of the 170 kilodalton and 180 kilodalton forms of topoisomerase II in NIH-3T3 cells. Cell Growth Differ 2, 209–214.

45. Kimura, K., Saijo, M., Ui, M., and Enomoto, T. (1994). Growth state- and cell cycle-dependent fluctuation in the expression of two forms of DNA topoisomerase II and possible specific modification of the higher molecular weight form in the M phase. J Biol Chem 269, 1173–1176.

46. Lee, J.H., and Berger, J.M. (2019). Cell Cycle-Dependent Control and Roles of DNA Topoisomerase II. Genes 10. 10.3390/genes10110859.

47. Hopfner, R., Mousli, M., Jeltsch, J.M., Voulgaris, A., Lutz, Y., Marin, C., Bellocq, J.P., Oudet, P., and Bronner, C. (2000). ICBP90, a novel human CCAAT binding protein, involved in the regulation of topoisomerase IIalpha expression. Cancer Res 60, 121–128.

48. Magan, N., Szremska, A.P., Isaacs, R.J., and Stowell, K.M. (2003). Modulation of DNA topoisomerase II alpha promoter activity by members of the Sp (specificity protein) and NF-Y (nuclear factor Y) families of transcription factors. Biochem J 374, 723–729. 10.1042/BJ20030032.

49. Liao, H., Winkfein, R.J., Mack, G., Rattner, J.B., and Yen, T.J. (1995). CENP-F is a protein of the nuclear matrix that assembles onto kinetochores at late G2 and is rapidly degraded after mitosis. J Cell Biol 130, 507–518. 10.1083/jcb.130.3.507.

50. Yang, Z.Y., Guo, J., Li, N., Qian, M., Wang, S.N., and Zhu, X.L. (2003). Mitosin/CENP-F is a conserved kinetochore protein subjected to cytoplasmic dynein-mediated poleward transport. Cell Res 13, 275–283. 10.1038/sj.cr.7290172.

51. Chan, G.K., Jablonski, S.A., Sudakin, V., Hittle, J.C., and Yen, T.J. (1999). Human BUBR1 is a mitotic checkpoint kinase that monitors CENP-E functions at kinetochores and binds the cyclosome/APC. J Cell Biol 146, 941–954. 10.1083/jcb.146.5.941.

52. Shin, H.J., Baek, K.H., Jeon, A.H., Park, M.T., Lee, S.J., Kang, C.M., Lee, H.S., Yoo, S.H., Chung, D.H., Sung, Y.C., et al. (2003). Dual roles of human BubR1, a mitotic checkpoint kinase, in the monitoring of chromosomal instability. Cancer Cell 4, 483–497. 10.1016/s1535-6108(03)00302-7.

53. Tatsumoto, T., Xie, X., Blumenthal, R., Okamoto, I., and Miki, T. (1999). Human Ect2 Is an Exchange Factor for Rho Gtpases, Phosphorylated in G2/M Phases, and Involved in Cytokinesis. Journal of Cell Biology 147, 921–928. 10.1083/jcb.147.5.921.

54. Liu, X., and Winey, M. (2012). The MPS1 Family of Protein Kinases. Annu Rev Biochem 81, 561–585. 10.1146/annurev-biochem-061611-090435.

55. Zhang, Y., Tan, L., Yang, Q., Li, C., and Liou, Y.-C. (2018). The microtubule-associated protein HURP recruits the centrosomal protein TACC3 to regulate K-fiber formation and support chromosome congression. Journal of Biological Chemistry 293, 15733–15747. 10.1074/jbc.RA118.003676.

56. Klosin, A., Oltsch, F., Harmon, T., Honigmann, A., Jülicher, F., Hyman, A.A., and Zechner, C. (2020). Phase separation provides a mechanism to reduce noise in cells. Science 367, 464–468. 10.1126/science.aav6691.

57. Azzam, M.E., and Algranati, I.D. (1973). Mechanism of Puromycin Action: Fate of Ribosomes after Release of Nascent Protein Chains from Polysomes. Proc. Natl. Acad. Sci. U.S.A. 70, 3866–3869. 10.1073/pnas.70.12.3866.

58. Fan, X.C., and Steitz, J.A. (1998). HNS, a nuclear-cytoplasmic shuttling sequence in HuR. Proc Natl Acad Sci U S A 95, 15293–15298. 10.1073/pnas.95.26.15293.

59. Wang, W., Caldwell, M.C., Lin, S., Furneaux, H., and Gorospe, M. (2000). HuR regulates cyclin A and cyclin B1 mRNA stability during cell proliferation. EMBO J 19, 2340–2350. 10.1093/emboj/19.10.2340.

60. Wu, X., Lan, L., Wilson, D.M., Marquez, R.T., Tsao, W.-C., Gao, P., Roy, A., Turner, B.A., McDonald, P., Tunge, J.A., et al. (2015). Identification and validation of novel small molecule disruptors of HuR-mRNA interaction. ACS Chem Biol 10, 1476–1484. 10.1021/cb500851u.

61. Frenkel-Morgenstern, M., Danon, T., Christian, T., Igarashi, T., Cohen, L., Hou, Y.-M., and Jensen, L.J. (2012). Genes adopt non-optimal codon usage to generate cell cycle-dependent oscillations in protein levels. Mol Syst Biol 8, 572. 10.1038/msb.2012.3.

62. Tanenbaum, M.E., Stern-Ginossar, N., Weissman, J.S., and Vale, R.D. (2015). Regulation of mRNA translation during mitosis. eLife 4, e07957. 10.7554/eLife.07957.

63. Cardona, A.H., Ecsedi, S., Khier, M., Yi, Z., Bahri, A., Ouertani, A., Valero, F., Labrosse, M., Rouquet, S., Robert, S., et al. (2023). Self-demixing of mRNA copies buffers mRNA:mRNA and mRNA:regulator stoichiometries. Cell 186, 4310–4324.e23. 10.1016/j.cell.2023.08.018.

64. Riback, J.A., Zhu, L., Ferrolino, M.C., Tolbert, M., Mitrea, D.M., Sanders, D.W., Wei, M.-T., Kriwacki, R.W., and Brangwynne, C.P. (2020). Composition-dependent thermodynamics of intracellular phase separation. Nature 581, 209–214. 10.1038/s41586-020-2256-2.

65. Sanchez-Burgos, I., Herriott, L., Collepardo-Guevara, R., and Espinosa, J.R. (2023). Surfactants or scaffolds? RNAs of different lengths exhibit heterogeneous distributions and play diverse roles in RNA-protein condensates. Biophysical Journal. 10.1016/j.bpj.2023.03.006.

66. Sanchez-Burgos, I., Espinosa, J.R., Joseph, J.A., and Collepardo-Guevara, R. (2022). RNA length has a non-trivial effect in the stability of biomolecular condensates formed by RNA-binding proteins. PLOS Computational Biology 18, e1009810. 10.1371/journal.pcbi.1009810.

67. Cochard, A., Garcia-Jove Navarro, M., Piroska, L., Kashida, S., Kress, M., Weil, D., and Gueroui, Z. (2022). RNA at the surface of phase-separated condensates impacts their size and number. Biophysical Journal 121, 1675–1690. 10.1016/j.bpj.2022.03.032.

68. Grant, G.D., Kedziora, K.M., Limas, J.C., Cook, J.G., and Purvis, J.E. (2018). Accurate delineation of cell cycle phase transitions in living cells with PIP-FUCCI. Cell Cycle 17, 2496–2516. 10.1080/15384101.2018.1547001.

69. Sakaue-Sawano, A., Kurokawa, H., Morimura, T., Hanyu, A., Hama, H., Osawa, H., Kashiwagi, S., Fukami, K., Miyata, T., Miyoshi, H., et al. (2008). Visualizing spatiotemporal dynamics of multicellular cell-cycle progression. Cell 132, 487–498. 10.1016/j.cell.2007.12.033.

70. Chouaib, R., Safieddine, A., Pichon, X., Imbert, A., Kwon, O.S., Samacoits, A., Traboulsi, A.-M., Robert, M.-C., Tsanov, N., Coleno, E., et al. (2020). A Dual Protein-mRNA Localization Screen Reveals Compartmentalized Translation and Widespread Co-translational RNA Targeting. Developmental Cell 54, 773–791.e5. 10.1016/j.devcel.2020.07.010.

71. Safieddine, A., Coleno, E., Salloum, S., Imbert, A., Traboulsi, A.-M., Kwon, O.S., Lionneton, F., Georget, V., Robert, M.-C., Gostan, T., et al. (2021). A choreography of centrosomal mRNAs reveals a conserved localization mechanism involving active polysome transport. Nat Commun 12, 1352. 10.1038/s41467-021-21585-7.

72. Dobin, A., Davis, C.A., Schlesinger, F., Drenkow, J., Zaleski, C., Jha, S., Batut, P., Chaisson, M., and Gingeras, T.R. (2013). STAR: ultrafast universal RNA-seq aligner. Bioinformatics 29, 15–21. 10.1093/bioinformatics/bts635.

73. Goecks, J., Nekrutenko, A., and Taylor, J. (2010). Galaxy: a comprehensive approach for supporting accessible, reproducible, and transparent computational research in the life sciences. Genome Biol 11, 1–13. 10.1186/gb-2010-11-8-r86.

74. Liao, Y., Smyth, G.K., and Shi, W. (2014). featureCounts: an efficient general purpose program for assigning sequence reads to genomic features. Bioinformatics 30, 923–930. 10.1093/bioinformatics/btt656.

75. Wu, T., Hu, E., Xu, S., Chen, M., Guo, P., Dai, Z., Feng, T., Zhou, L., Tang, W., Zhan, L., et al. (2021). clusterProfiler 4.0: A universal enrichment tool for interpreting omics data. Innovation 2. 10.1016/j.xinn.2021.100141.

76. Yu, G., Wang, L.-G., Han, Y., and He, Q.-Y. (2012). clusterProfiler: an R Package for Comparing Biological Themes Among Gene Clusters. OMICS: A Journal of Integrative Biology 16, 284–287. 10.1089/omi.2011.0118.

77. Safieddine, A., Coleno, E., Lionneton, F., Traboulsi, A.-M., Salloum, S., Lecellier, C.-H., Gostan, T., Georget, V., Hassen-Khodja, C., Imbert, A., et al. (2023). HT-smFISH: a cost-effective and flexible workflow for high-throughput single-molecule RNA imaging. Nat Protoc 18, 157–187. 10.1038/s41596-022-00750-2.

78. Schindelin, J., Arganda-Carreras, I., Frise, E., Kaynig, V., Longair, M., Pietzsch, T., Preibisch, S., Rueden, C., Saalfeld, S., Schmid, B., et al. (2012). Fiji: an open-source platform for biological-image analysis. Nat Methods 9, 676–682. 10.1038/nmeth.2019.

79. Allan, C., Burel, J.-M., Moore, J., Blackburn, C., Linkert, M., Loynton, S., MacDonald, D., Moore, W.J., Neves, C., Patterson, A., et al. (2012). OMERO: flexible, model-driven data management for experimental biology. Nat Methods 9, 245–253. 10.1038/nmeth.1896.

80. Stringer, C., Wang, T., Michaelos, M., and Pachitariu, M. (2021). Cellpose: a generalist algorithm for cellular segmentation. Nat Methods 18, 100–106. 10.1038/s41592-020-01018-x.

81. Imbert, A., Ouyang, W., Safieddine, A., Coleno, E., Zimmer, C., Bertrand, E., Walter, T., and Mueller, F. (2022). FISH-quant v2: a scalable and modular tool for smFISH image analysis. RNA 28, 786–795. 10.1261/rna.079073.121.

82. Mueller, F., Senecal, A., Tantale, K., Marie-Nelly, H., Ly, N., Collin, O., Basyuk, E., Bertrand, E., Darzacq, X., and Zimmer, C. (2013). FISH-quant: automatic counting of transcripts in 3D FISH images. Nat. Methods 10, 277–278. 10.1038/nmeth.2406.

83. Balak, C., Benard, M., Schaefer, E., Iqbal, S., Ramsey, K., Ernoult-Lange, M., Mattioli, F., Llaci, L., Geoffroy, V., Courel, M., et al. (2019). Rare De Novo Missense Variants in RNA Helicase DDX6 Cause Intellectual Disability and Dysmorphic Features and Lead to P-Body Defects and RNA Dysregulation. Am J Hum Genet 105, 509–525. 10.1016/j.ajhg.2019.07.01

