## Supplemental figures for "Cell cycle-dependent mRNA localization in P-bodies"

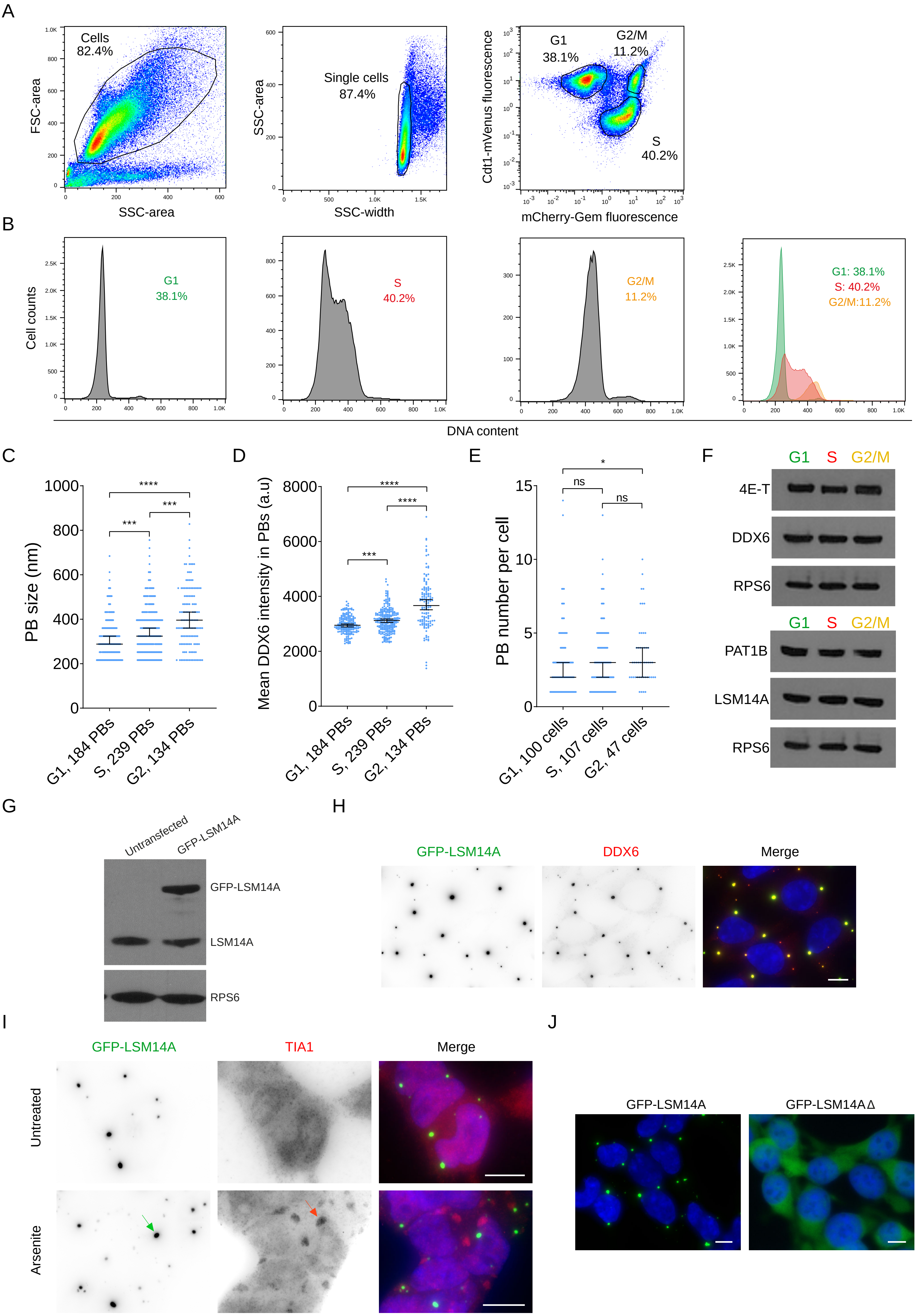

A
Cells
82.4%
FSC-area
SSC-area
Single cells
87.4%
SSC-area
SSC-width
G1
38.1%
G2/M
11.2%
S
40.2%
Cdt1-mVenus fluorescence
mCherry-Gem fluorescence
B
Cell counts
G1
38.1%
S
40.2%
G2/M
11.2%
G1: 38.1%
S: 40.2%
G2/M:11.2%
DNA content
C
PB size (nm)
G1, 184 PBs
S, 239 PBs
G2, 134 PBs
D
Mean DDX6 intensity in PBs (a.u)
G1, 184 PBs
S, 239 PBs
G2, 134 PBs
E
PB number per cell
ns
ns
G1, 100 cells
S, 107 cells
G2, 47 cells
F
G1 S G2/M
4E-T
DDX6
RPS6
G1 S G2/M
PAT1B
LSM14A
RPS6
G
Untransfected
GFP-LSM14A
GFP-LSM14A
LSM14A
RPS6
H
GFP-LSM14A
DDX6
Merge
I
GFP-LSM14A
TIA1
Merge
Untreated
Arsenite
J
GFP-LSM14A
GFP-LSM14AΔ

**Figure S1: Characterization and benchmarking of HEK293-FUCCI and HEK293 GFP-LSM14A cells, related to Figure 1**. (A) Gating strategy based on cell size (left panel), restriction to single cells (middle panel) and PIP-FUCCI signal (right panel). (B) Histograms of Hoechst intensity (DNA content) for each of the three populations gated in (A) (G1, S, and G2/M). The merged three panels are presented below. (C-E) Distribution of PB size (C), mean DDX6 intensity in PBs (D), and PB number per cell (E) across the cell cycle. Horizontal line, median; error bars, 95% CI. The numbers of PBs or cells are indicated on the x-axis (n=2 experiments). Two-tailed Mann-Whitney tests: ****, $p<0.0001$; ***, $p<0.001$; *, $p<0.05$; ns, non-significant ($p>0.05$). (F) Western blot analysis of cytoplasmic levels of several PB proteins across the cell cycle. The ribosomal protein RPS6 was used as a loading control. (G) Western blot analysis of LSM14A in untransfected HEK293 cells and in the clone used for PB purification by FAPS. The ribosomal protein RPS6 was used as a loading control. (H) Widefield fluorescence microscopy images of HEK293 cells expressing GFP-LSM14A (left panel, in green in the merge), immunostained with an anti-DDX6 antibody (middle panel, in red in the merge). Nuclei were stained with DAPI (in blue). Scale bar, 10 µm. (I) Same as H after immunostaining of the stress granule marker TIA1, in cells either untreated (upper panel) or treated for 30 min with arsenite (lower panel). The green and red arrows point to a PB and a stress granule, respectively. (J) Widefield images of cells expressing GFP-LSM14A or a truncated version of GFP-LSM14A that does not localize in PBs (in green). Nuclei were stained with DAPI (in blue). Scale bars, 10 µm.

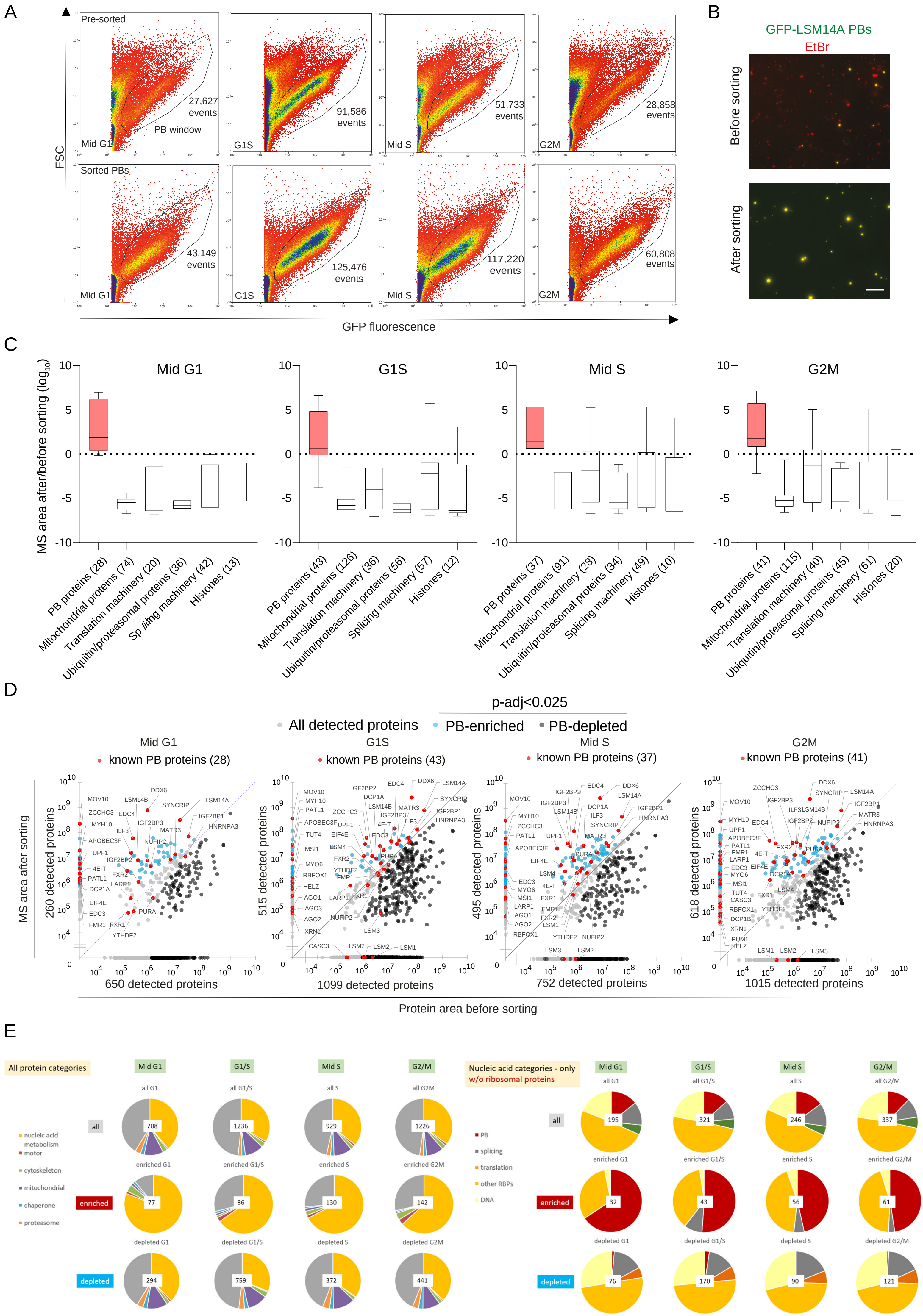

A
FSC
GFP fluorescence
Pre-sorted
Sorted PBs
Mid G1
G1S
Mid S
G2M
PB window
27,627 events
91,586 events
51,733 events
28,858 events
43,149 events
125,476 events
117,220 events
60,808 events
B
GFP-LSM14A PBs
EtBr
Before sorting
After sorting
C
MS area after/before sorting (log10)
Mid G1
G1S
Mid S
G2M
PB proteins (28)
Mitochondrial proteins (74)
Translation machinery (20)
Ubiquitin/proteasomal proteins (36)
Splicing machinery (42)
Histones (13)
PB proteins (43)
Mitochondrial proteins (126)
Translation machinery (36)
Ubiquitin/proteasomal proteins (56)
Splicing machinery (57)
Histones (12)
PB proteins (37)
Mitochondrial proteins (91)
Translation machinery (28)
Ubiquitin/proteasomal proteins (34)
Splicing machinery (49)
Histones (10)
PB proteins (41)
Mitochondrial proteins (115)
Translation machinery (40)
Ubiquitin/proteasomal proteins (45)
Splicing machinery (61)
Histones (20)
D
p-adj<0.025
All detected proteins
PB-enriched
PB-depleted
known PB proteins (28)
known PB proteins (43)
known PB proteins (37)
known PB proteins (41)
MS area after sorting
260 detected proteins
515 detected proteins
495 detected proteins
618 detected proteins
650 detected proteins
1099 detected proteins
752 detected proteins
1015 detected proteins
Protein area before sorting
E
All protein categories
Mid G1
G1/S
Mid S
G2/M
all
enriched
depleted
nucleic acid metabolism
motor
cytoskeleton
mitochondrial
chaperone
proteasome
708
1236
929
1226
77
86
130
142
294
759
372
441
Nucleic acid categories - only w/o ribosomal proteins
PB
splicing
translation
other RBPs
DNA
195
321
246
337
32
43
56
61
76
170
90
121

**Figure S2: FAPS and MS analysis before and after PB purification across the cell cycle, related to Figure 1.** (A) Representative FAPS profiles are shown for mid G1, G1S, mid S and G2M cells before (upper panels) and after (lower panels) sorting, as in Figure 1E. The Figure 1E right panels corresponding to G1S were included to facilitate comparison. (B) Widefield fluorescence images of lysates before and after sorting. PBs were labelled with GFP-LSM14A (green) and contaminants were revealed by non-specific ethidium bromide (EtBr) staining (red). Scale bar, 10 µm. (C) The data are presented as in Figure 1G. The Figure 1G panel corresponding to G1S was included to facilitate comparison. (D) Scatter plots highlighting detected proteins before and after sorting across the cell cycle. Grey dots correspond to all detected proteins (protein area >1) while blue and black ones correspond to PB-enriched or depleted proteins respectively (p-adj<0.025 based on a Fisher test). Known PB proteins are shown in red, including FXR1 and FXR2 confirmed in Figure 1 (list in Table S1). The diagonal is in purple. (E) Pie charts showing the fractions of different categories of PB enriched or depleted proteins, with the nucleic acid metabolism category detailed on the right.

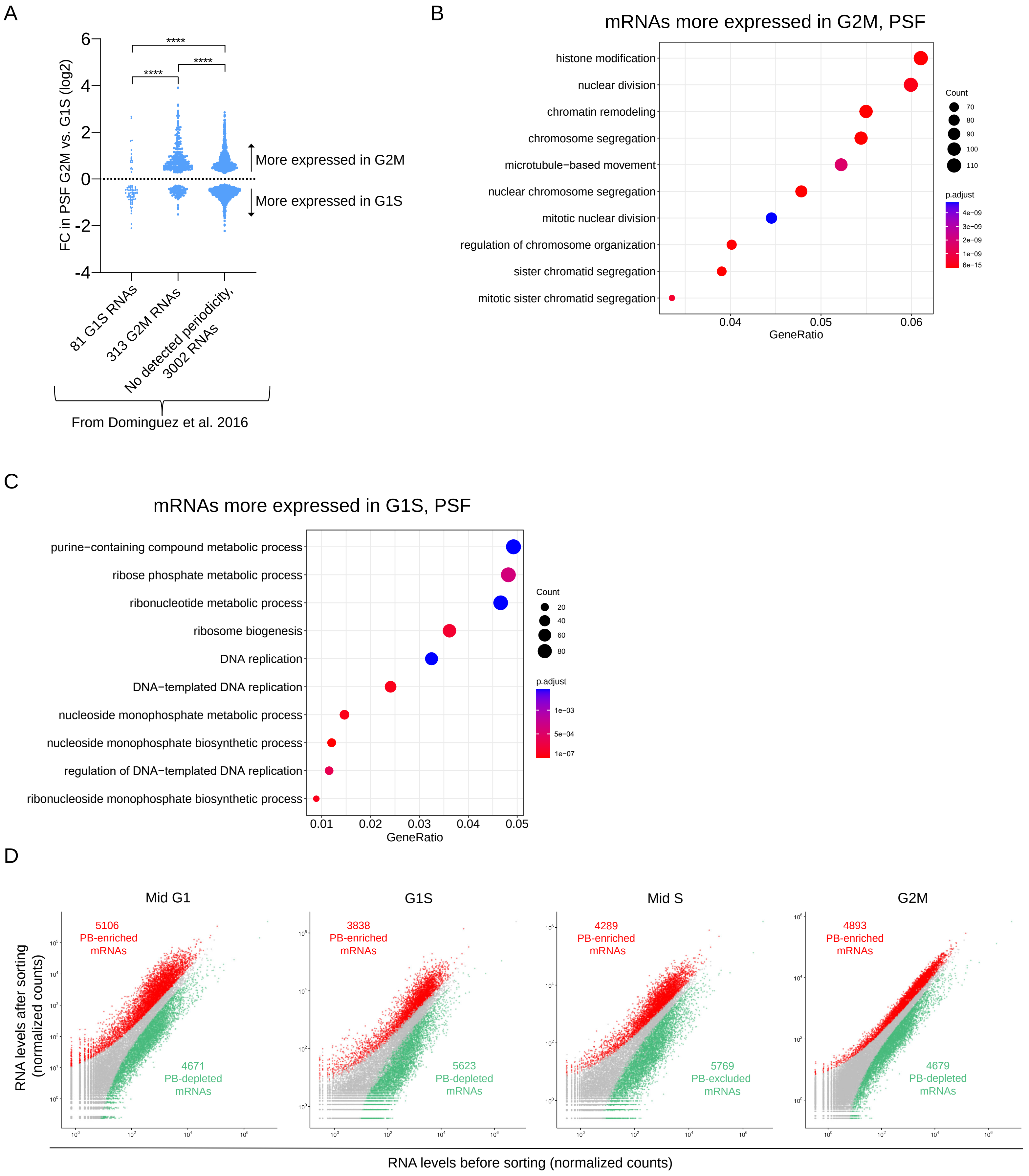

A
FC in PSF G2M vs. G1S (log2)
6
4
2
0
-2
-4
****
****
****
More expressed in G2M
More expressed in G1S
81 G1S RNAs
313 G2M RNAs
No detected periodicity, 3002 RNAs
From Dominguez et al. 2016
B
mRNAs more expressed in G2M, PSF
histone modification
nuclear division
chromatin remodeling
chromosome segregation
microtubule−based movement
nuclear chromosome segregation
mitotic nuclear division
regulation of chromosome organization
sister chromatid segregation
mitotic sister chromatid segregation
0.04
0.05
0.06
GeneRatio
Count
70
80
90
100
110
p.adjust
4e−09
3e−09
2e−09
1e−09
6e−15
C
mRNAs more expressed in G1S, PSF
purine−containing compound metabolic process
ribose phosphate metabolic process
ribonucleotide metabolic process
ribosome biogenesis
DNA replication
DNA−templated DNA replication
nucleoside monophosphate metabolic process
nucleoside monophosphate biosynthetic process
regulation of DNA−templated DNA replication
ribonucleoside monophosphate biosynthetic process
0.01
0.02
0.03
0.04
0.05
GeneRatio
Count
20
40
60
80
p.adjust
1e−03
5e−04
1e−07
D
Mid G1
5106 PB-enriched mRNAs
4671 PB-depleted mRNAs
G1S
3838 PB-enriched mRNAs
5623 PB-depleted mRNAs
Mid S
4289 PB-enriched mRNAs
5769 PB-excluded mRNAs
G2M
4893 PB-enriched mRNAs
4679 PB-depleted mRNAs
RNA levels after sorting (normalized counts)
RNA levels before sorting (normalized counts)

**Figure S3: RNA changes between G1S and G2M in the pre-sorting fraction and RNA content before and after PB purification within each cell cycle phase, related to Figure 2.** (A) Fold changes observed between G1S and G2M in the pre-sorting fractions (PSF) for RNAs identified as cyclic or not by Dominguez et al, 2016[30]. Only well-detected (average normalized counts>100) and significant fold changes (p-adj<0.05) are shown. Two-tailed Mann-Whitney tests: ****, p<0.0001. (B) GO analyses of mRNAs that are significantly more expressed (p-adj<0.05) in G2M than in G1S before sorting. Representation is as in Fig 2E. (C) Same as B for mRNAs that are significantly more expressed in G1S than in G2M. (D) Comparison of RNA levels (in normalized counts) before and after sorting, as in Figure 2A. The Figure 2A panel corresponding to mid S cells was included to facilitate comparison.

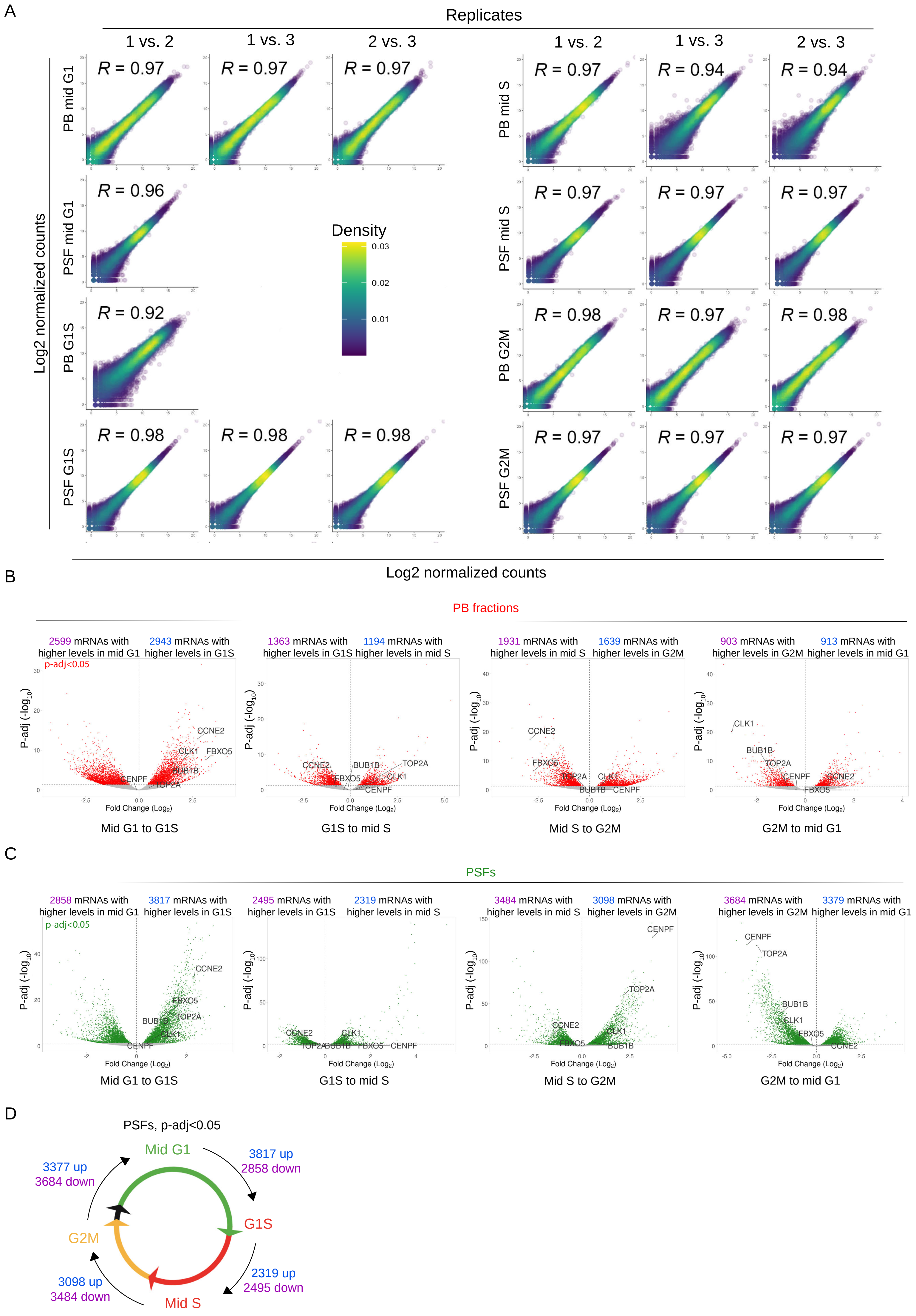
A
Replicates
1 vs. 2
1 vs. 3
2 vs. 3
PB mid G1
PSF mid G1
PB G1S
PSF G1S
PB mid S
PSF mid S
PB G2M
PSF G2M
Log2 normalized counts
Density
R = 0.97
R = 0.96
R = 0.92
R = 0.98
R = 0.94
B
PB fractions
2599 mRNAs with higher levels in mid G1
2943 mRNAs with higher levels in G1S
p-adj<0.05
1363 mRNAs with higher levels in G1S
1194 mRNAs with higher levels in mid S
1931 mRNAs with higher levels in mid S
1639 mRNAs with higher levels in G2M
903 mRNAs with higher levels in G2M
913 mRNAs with higher levels in mid G1
P-adj (-log10)
Fold Change (Log2)
CCNE2
CLK1
FBXO5
BUB1B
CENPF
TOP2A
Mid G1 to G1S
G1S to mid S
Mid S to G2M
G2M to mid G1
C
PSFs
2858 mRNAs with higher levels in mid G1
3817 mRNAs with higher levels in G1S
p-adj<0.05
2495 mRNAs with higher levels in G1S
2319 mRNAs with higher levels in mid S
3484 mRNAs with higher levels in mid S
3098 mRNAs with higher levels in G2M
3684 mRNAs with higher levels in G2M
3379 mRNAs with higher levels in mid G1
Mid G1 to G1S
G1S to mid S
Mid S to G2M
G2M to mid G1
D
PSFs, p-adj<0.05
Mid G1
G1S
Mid S
G2M
3817 up
2858 down
2319 up
2495 down
3098 up
3484 down
3377 up
3684 down

**Figure S4: Pairwise correlation between RNA-seq replicates and the evolution of the RNA content before and after sorting between successive cell cycle phases, related to Figure 2.** (A) Comparison of RNA levels (in normalized counts) within replicates before and after sorting. The Pearson correlation coefficient (R) is indicated. All experiments were performed in triplicates. One replicate before sorting (from mid G1) and one after sorting (from G1S) were removed due to $R<0.9$, leaving 2 replicates for these conditions. (B) Volcano plots showing the changes in mRNAs between successive cell cycle phases in purified PBs. The representation is as in Figure 2C. (C) Same as A, but in the PSF. (D) Schematic summary of significant changes (p-adj<0.05) across the cell cycle in the PSF.

A

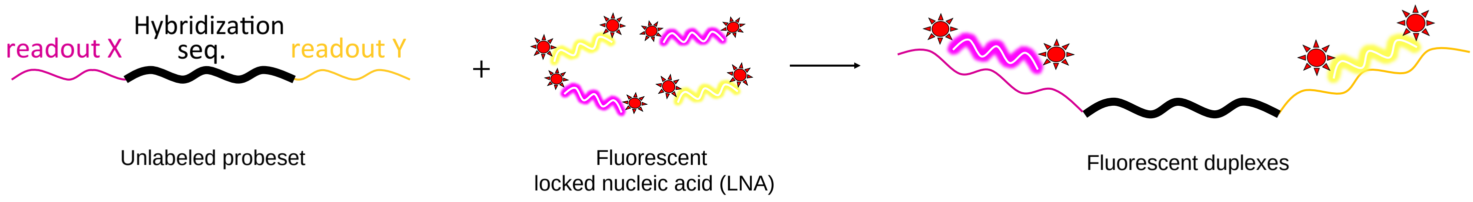


B

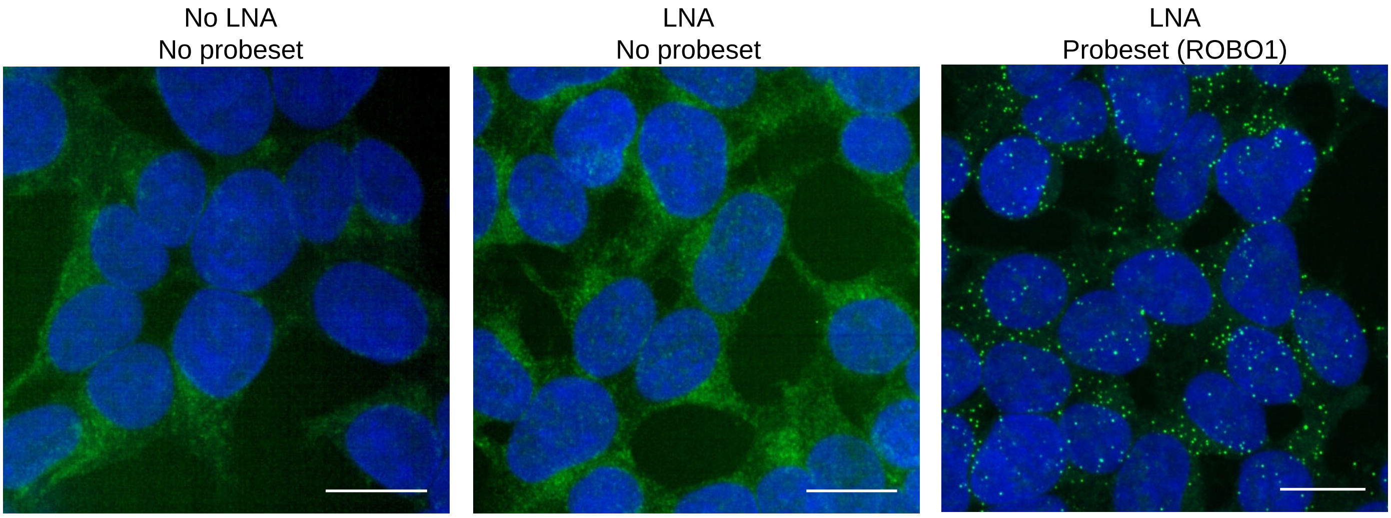


C

| mRNA | G1 cell number | G2 cell number |
|---|---|---|
| TTK | 426 | 140 |
| ANLN | 343 | 113 |
| DEPDC1 | 810 | 267 |
| ECT2 | 666 | 219 |
| DLGAP5 | 596 | 198 |
| MKI67 | 476 | 157 |
| ROBO1 | 524 | 173 |
| GPBP1 | 298 | 96 |

**Figure S5: Technical controls and cell numbers for HT-smFISH, related to Figure 5.** (A) Principle of HT-smFISH probe design. Primary unlabeled probesets contain a mRNA specific sequence, flanked by 2 readout sequences that in turn bind fluorescently-labeled locked nucleid acids (LNAs). Described in detail in Safieddine et al.[77] (B) Routine negative controls used in the screen. The mRNA channel is in green and DAPI stained nuclei in blue. Scale bars, 10 µm. (C) Table showing the number of cell analyzed and classified as either G1 or G2 for the mRNAs presented in Figure 5. The cell counts for remaining mRNAs are found in Supplementary Table 7.

**Figure S6: Differential mRNA localization does not depend on non-polysomal mRNA levels, related to Figure 6.** (A) Fraction of mRNAs localized in PBs as a function of their cytoplasmic expression levels, from smFISH experiments presented in Figures 6 (TOP2A) and 3 (FBXO5, CLK1). Each dot corresponds to one cell. The Spearman correlation coefficient are indicated. (B) TOP2A smFISH in HEK293-FUCCI cells transiently expressing LSM14A-Halo to label PBs in far-red, treated or not with puromycin for 1 hr. Left panel: cytoplasmic TOP2A mRNAs and nuclear mCherry-Gem signal (in red in the merge). Middle panel: Cdt1-mVenus (in green in the merge). Cells in early G1 were classified based on the combination of TOP2A mRNA and FUCCI labeling patterns, and cells in G2 based on the FUCCI system. In the merge, PBs are in blue and DAPI-stained nuclei in white. Scale bars, 10 and 1 µm in the main images and insets, respectively. (C) Metaphase HeLa cells expressing ASPM-MS2 mRNA, treated or not with puromycin for 15 min. The ASPM-MS2 mRNA was detected by smFISH against the MS2 sequence (Cy3, shown in green). Arrows indicate the typical ASPM mRNA accumulation on metaphase centrosomes. DAPI-stained DNA is in blue. Scale bars, 10 µm. (D) The enrichment of 4E-T targets with or without HuR binding sites or AREs in the top 1000 PB mRNAs across the cell cycle.

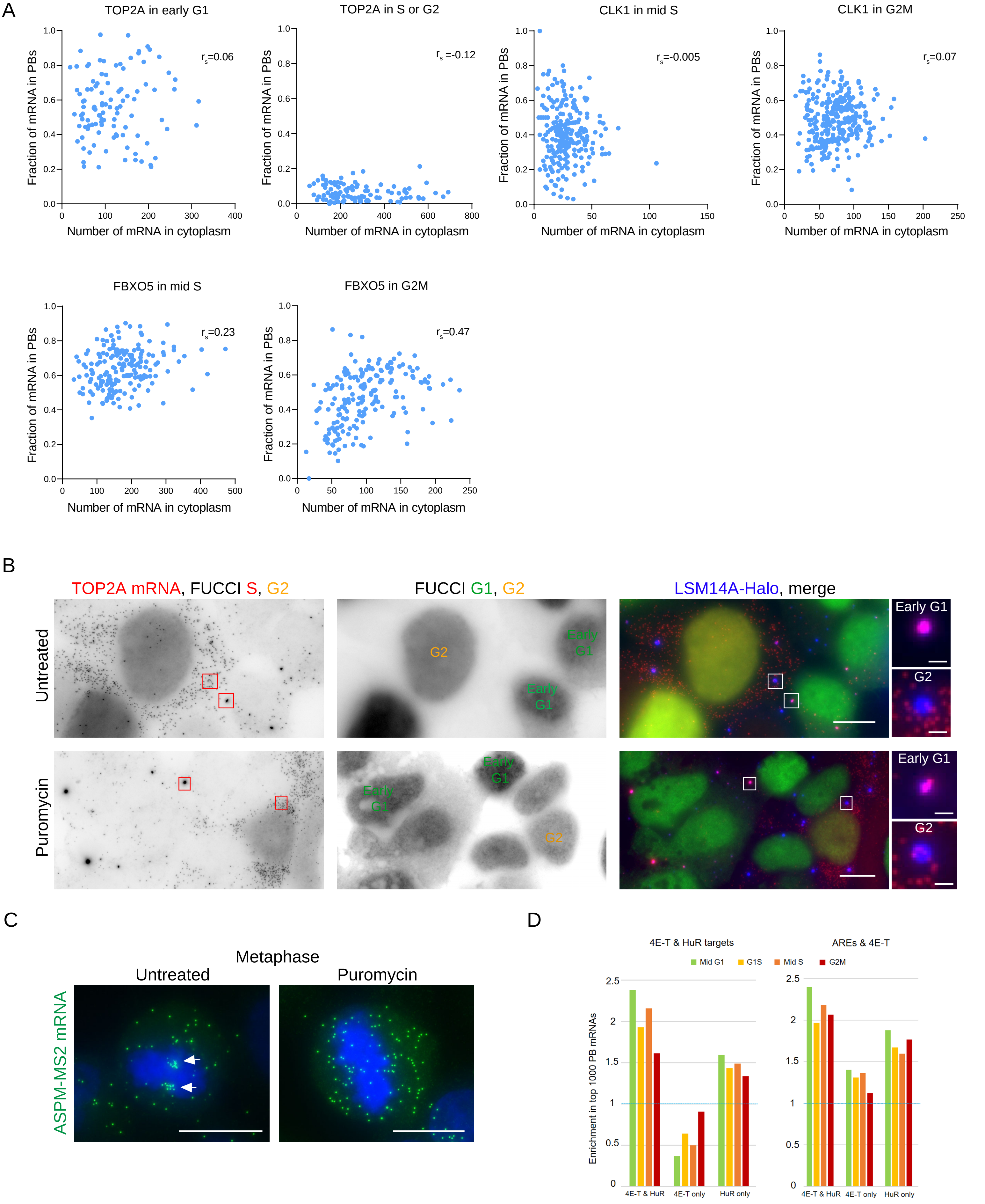
A
TOP2A in early G1
TOP2A in S or G2
CLK1 in mid S
CLK1 in G2M
FBXO5 in mid S
FBXO5 in G2M
Fraction of mRNA in PBs
Number of mRNA in cytoplasm
r_s=0.06
r_s =-0.12
r_s=-0.005
r_s=0.07
r_s=0.23
r_s=0.47
B
TOP2A mRNA, FUCCI S, G2
FUCCI G1, G2
LSM14A-Halo, merge
Untreated
Puromycin
Early G1
G2
C
Metaphase
Untreated
Puromycin
ASPM-MS2 mRNA
D
4E-T & HuR targets
AREs & 4E-T
Mid G1
G1S
Mid S
G2M
Enrichment in top 1000 PB mRNAs
4E-T & HuR
4E-T only
HuR only

A

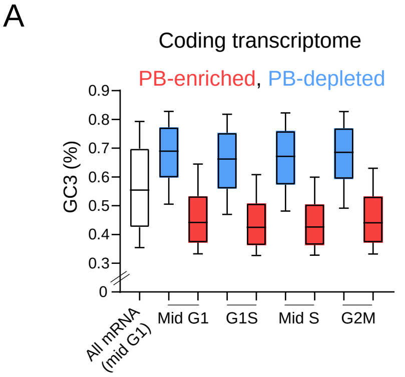


B

Correlation between PB-enrichment and GC content

| | Mid G1 | G1S | Mid S | G2M |
|---|---|---|---|---|
| GC, full mRNA | -0.78 | -0.79 | -0.79 | -0.75 |
| GC3 | -0.66 | -0.67 | -0.68 | -0.63 |
| 3'UTR | -0.61 | -0.63 | -0.62 | -0.59 |
| 5'UTR | -0.22 | -0.22 | -0.23 | -0.20 |

C

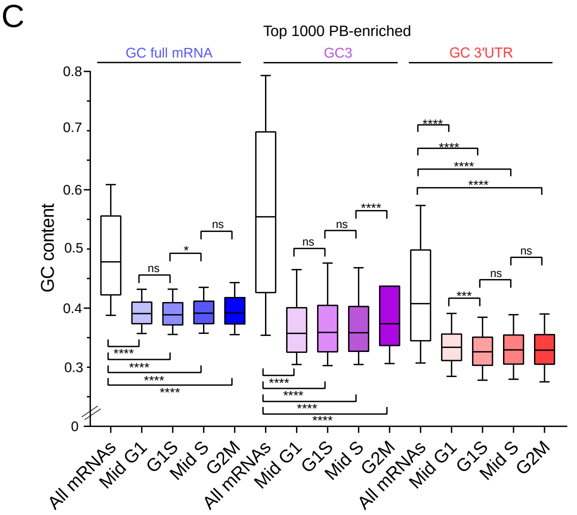


D

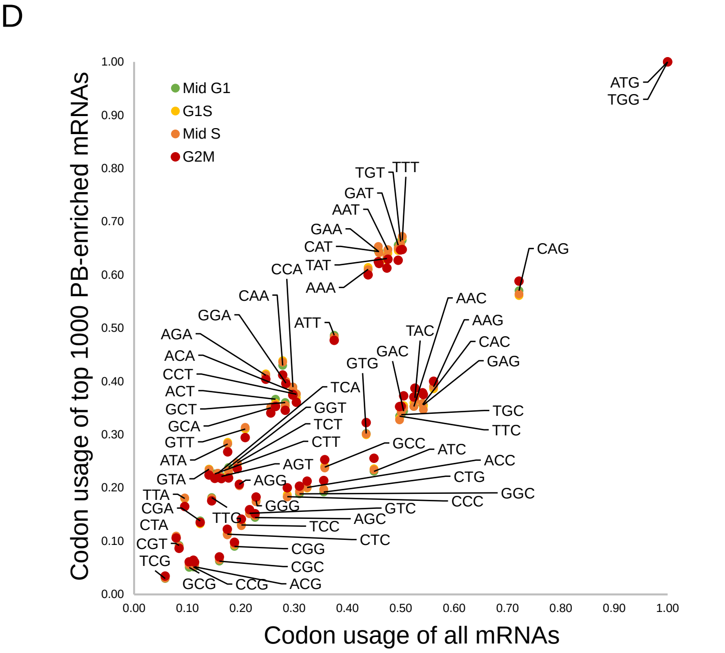


E

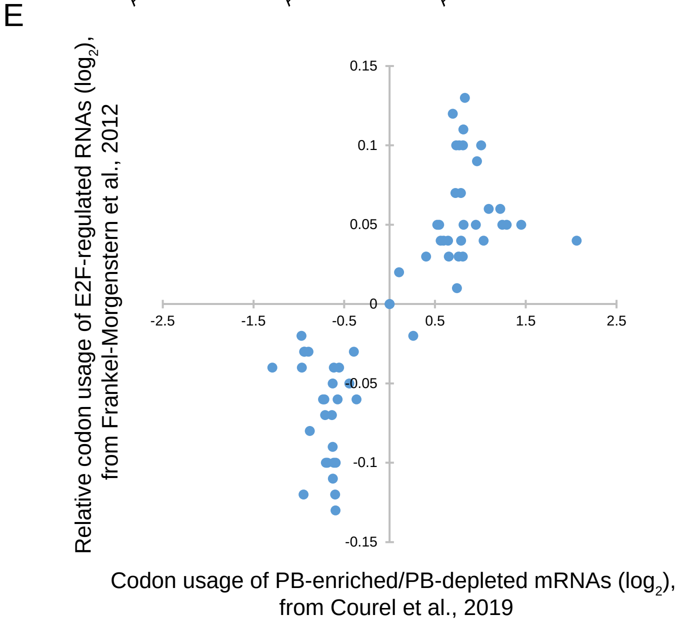


F

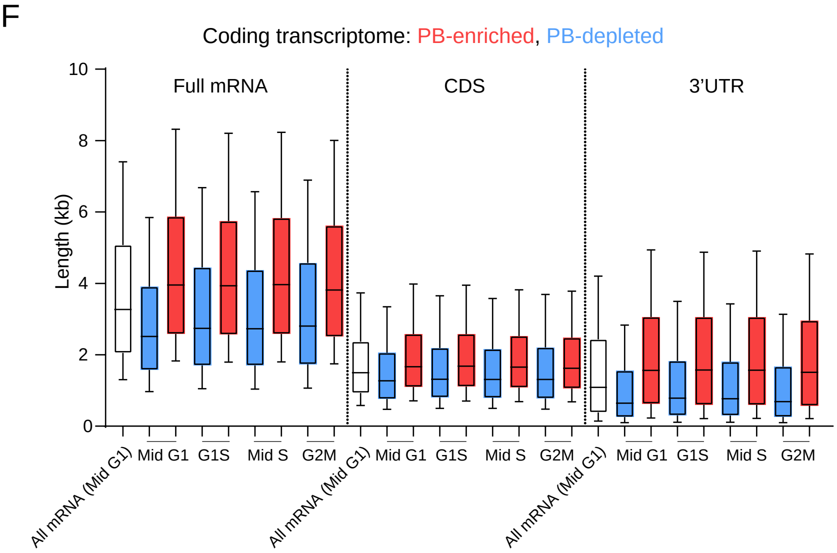


G

Correlation between PB-enrichment and mRNA length

| | Mid G1 | G1S | Mid S | G2M |
|---|---|---|---|---|
| Full mRNA | 0.33 | 0.22 | 0.24 | 0.2 |
| CDS | 0.18 | 0.07 | 0.1 | 0.04 |
| 3'UTR | 0.3 | 0.23 | 0.24 | 0.24 |

H

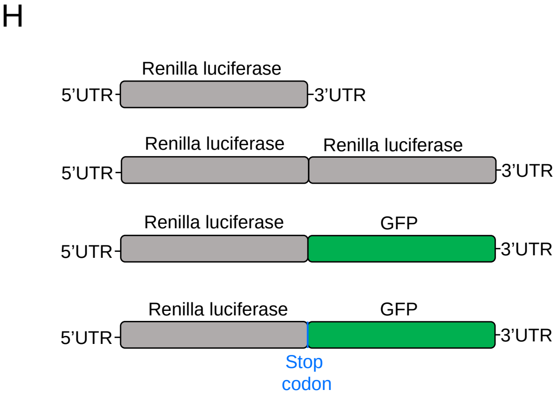


I

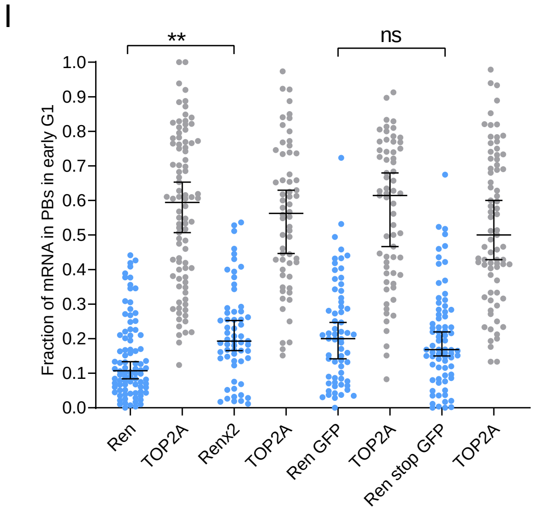

**Figure S7: Features of PB mRNAs across the cell cycle, related to Figure 7.** (A) GC content in the third codon position (GC3) for all detected mRNAs (normalized counts >100) (in white), or all PB-enriched (in red) or depleted (in blue) mRNAs (FC>0 or <0 respectively, p-adj<0.05, normalized counts >100) across the cell cycle. Whiskers represent the 10 to 90% percentile. (B) The Spearman correlation coefficients between GC contents and PB-enrichment for all expressed mRNAs (normalized counts >100). (C) GC content of the full mRNA, CDS, and 3'UTR for all detected mRNAs (all mRNAs, normalized counts >100) or for the top 1000 PB-enriched mRNAs (FC>0, p-adj<0.05, normalized counts >100) in the various cell cycle phases. Two-tailed Mann-Whitney tests: ****, p<0.0001; **, p<0.005; *, p<0.05; ns, non-significant (p>0.05). (D) Comparison of the codon usage frequency of all expressed mRNAs (normalized counts >100) with the codon usage of the top 1000 PB-enriched mRNAs (FC>0, p-adj<0.05, normalized counts >100) in the various cell cycle phases. (E) Comparison of the relative codon usage in PB-enriched vs. PB-depleted mRNAs in asynchronous cells (as reported by Courel et al., 2019[8]) with the relative codon usage in a group of E2F-induced cell cycle genes (as reported by Morgenstern et al., 2012[62]). (F) Length of the full mRNA, CDS, or 3'UTR of all detected mRNAs (normalized counts >100), or all PB-enriched or depleted mRNAs (FC>0 or <0 respectively, p-adj<0.05, normalized counts>100) across the cell cycle, represented as in (A). (G) The Spearman correlation coefficients between mRNA length and PB-enrichment for all expressed mRNAs (normalized counts >100). (H) Representation of various Renilla reporters. (I) The fraction of mRNA in PBs in early G1 cells across several Renilla reporters.
